# Temporal Shifts in Gene Expression Drive Quantitative Resistance to a Necrotrophic Fungus in a Tomato Crop Wild Relative

**DOI:** 10.1101/2025.07.17.665297

**Authors:** S. Einspanier, A. Tobian Herreno, N. J. Rangegowda, L. Groenenberg, T. van der Lee, R. Stam

## Abstract

Resistance breeding against generalist necrotrophic pathogens heavily relies on quantitative disease resistance (QDR). Lesion growth dynamics involve distinct phases (e.g., lag phase duration or lesion doubling time), each independently affecting the overall symptom severity. While the genetic and regulatory basis of lesion growth rate has been studied, the host-derived regulation of the lag-phase duration remains largely uncharacterised.

In this study, we tested the regulatory response of *Solanum pennellii* genotypes exhibiting different lag-phase durations. We conducted a time-series gene expression profiling experiment dissecting genotype-specific regulatory responses to *Sclerotinia sclerotiorum* inoculation. We observed genotype-specific regulatory trajectories, with resistant plants displaying early activation of defence-related genes during the asymptomatic phase. These genes, regulated by a WRKY6-centered gene regulatory network, exhibited elevated basal expression in resistant genotypes and a fine-tuned longitudinal expression with induction before lesion onset. In contrast, susceptible genotypes lacked this early response, showing gene induction only post-infection. This study is the first to link host regulatory dynamics to lag-phase duration, suggesting that elevated basal expression of receptor genes and a WRKY6-mediated gene regulatory network may enhance QDR. These findings provide insights into the regulatory foundation of QDR and establish a functional basis for more focused breeding of QDR traits.

## Background

Inoculating plant leaves with a necrotrophic pathogen, like *Sclerotinia sclerotiorum,* does not always cause instant lesion onset. This phase, marked by the absence of symptoms and fungal growth, is commonly referred to as the lag phase. Originally, the lag phase was mostly studied in human medicine, focusing on bacterial infectious diseases such as *Streptococcus pneumoniae* or *Escherichia coli*. This research led to the identification of microbial mechanisms determining the lag phase duration, such as microbial promoter activity or the structure of the polysaccharide capsule protecting bacteria (Madar et al. 2013; Hathaway et al. 2012). Although well understood in microbiology, modern plant pathology lacks a fundamental understanding of the processes determining this complex phenotype. Yet, the lag phase has recently gained attention in light of quantitative disease resistance (QDR).

During the last decades, the research on resistance breeding has mostly focused on investigating R-gene-mediated qualitative disease resistance (Chen et al. 2024; Kaur et al. 2021). Prominent examples like the tomato R gene *Pto,* which confers complete race-specific resistance against the bacterial pathogen *Pseudomonas syringae* pv. tomato, show the powerful capabilities of R-genes and have led to wide use in crops (Tang et al. 1999). However, R-genes against major necrotrophic pathogens like the cosmopolitan ascomycetes *Botrytis cinerea* and *S. sclerotiorum* are yet to be found (Bi et al. 2023; Derbyshire et al. 2022). Additionally, R-genes provide a strong selection pressure on pathogen populations and might lead to rapid pathogen adaptation and the loss of efficacy even when exploited as building blocks in R-gene pyramiding strategies (Stam and McDonald 2018). Thus, the polygenic QDR serves as an essential source of resistance traits (Roux et al. 2014; Poland et al. 2009; Corwin and Kliebenstein 2017).

QDR is a non-complete form of resistance (sometimes called disease tolerance) underlying complex polymorphic control (regulatorily and genomically). Targeted breeding for QDR presents major challenges because of the complexity of linking nuanced phenotypes with interconnected regulatory and genomic determinants (Roux et al. 2014; Derbyshire et al. 2022). Most QDR studies rely on single-time-point measurements of severity, which is often estimated by lesion size or area under the disease-progress curve (AUDPC) as the primary resistance phenotype. However, focusing solely on lesion severity might undervalue temporal components of disease progression, such as lag phase duration and lesion doubling time (LDT), which can be crucial elements of a plant’s successful QDR, as delayed infection or reduced growth speed might allow the host plant to complete its life cycle without large fitness costs.

We recently hypothesised that distinct parameters of the lesion growth dynamics, including the lag phase duration and LDT, are host-dependent QDR phenotypes, each contributing independently to symptom severity in a genotype-specific manner (Einspanier et al. 2024). Defining these two metrics as separate drivers of QDR allows for the functional characterisation of independent resistance mechanisms. Here, we focus on lag phase duration, investigating its role in shaping quantitative resistance and its potential as a tool in breeding for QDR.

Most research in both humans and plants indicates that the duration of the lag phase is primarily influenced by the pathogen’s genotype (Barbacci et al. 2020; Hamill et al. 2020). Accordingly, several reports indicated that infection delays caused by necrotrophic pathogens are primarily associated with fungal characteristics like secreted proteins (effectors) and phytotoxins (Leisen et al. 2022; Denton-Giles et al. 2020). However, Hamill et al. (2020) demonstrated that cellular stress in microbial model organisms can significantly alter lag phase duration, which, in turn, may be influenced by the host genotype. Similarly, in a previous study on tomato crop wild relatives, we found that lag-phase duration varies significantly across different host genotypes (Einspanier et al. 2024). This observation aligns with recent reports by Sperfeld et al. (2024), who showed that methylated compounds secreted by photosynthetic organisms can alter the lag phase duration of the bacteria *Phaeobacter inhibens*. It is crucial to emphasise that we have shown a strong connection between delayed lesion onset of *S. sclerotiorum* infection and fungal development, distinguishing it from an asymptomatic growth phase, often referred to as the latency phase (Einspanier et al. 2024).

Building on recent insights, we hypothesise that host-specific regulatory mechanisms, besides pre-formed morphological properties, influence the lag-phase-mediated level of QDR against *S. sclerotiorum*. We speculate that early host-parasite interactions play a critical role in disease progression, possibly involving a complex network of genes responsible for detoxification, phytoalexin transport, and stress response. To assess early stress responses, we use the maximum quantum efficiency of PSII (Fv/Fm) as a key physiological marker.

We further hypothesise that the lag phase duration is controlled by a host-specific regulatory pattern, where gene expression composition, orchestration, and magnitude ultimately determine the disease outcome. To test this, we conducted RNA sequencing on a time-series experiment integrated with high-resolution phenotyping. By linking RGB-imagery and Fv/Fm measurements to longitudinal gene regulatory clusters and gene regulatory networks, we provide new insights into the determinants of lag-phase duration as a key factor of QDR.

The findings of this study leverage our understanding of QDR and its pleiotropic regulation, with clear applications in crop improvement for durable disease resistance.

## Results

### *S. pennellii* genotypes harbour significant QDR diversity

In a previous study, we performed a high-resolution phenotyping experiment to characterise the lag phase duration on different Solanum species using commonly available RGB cameras (Einspanier et al. 2024). We observed significant differences in the lag phase duration between the *S. pennellii* accessions LA1809, LA1941 and LA1282. We measured the shortest lag phase duration on the genotype LA1809 (lsmean 1.59d, 38.16 h), an intermediary lag phase duration of 2.08d (49.92 h) on LA1941, and a significantly increased lag phase duration on LA1282 (lsmean 2.37 d, 56.88h, fig. 1A).

**Figure 1:**
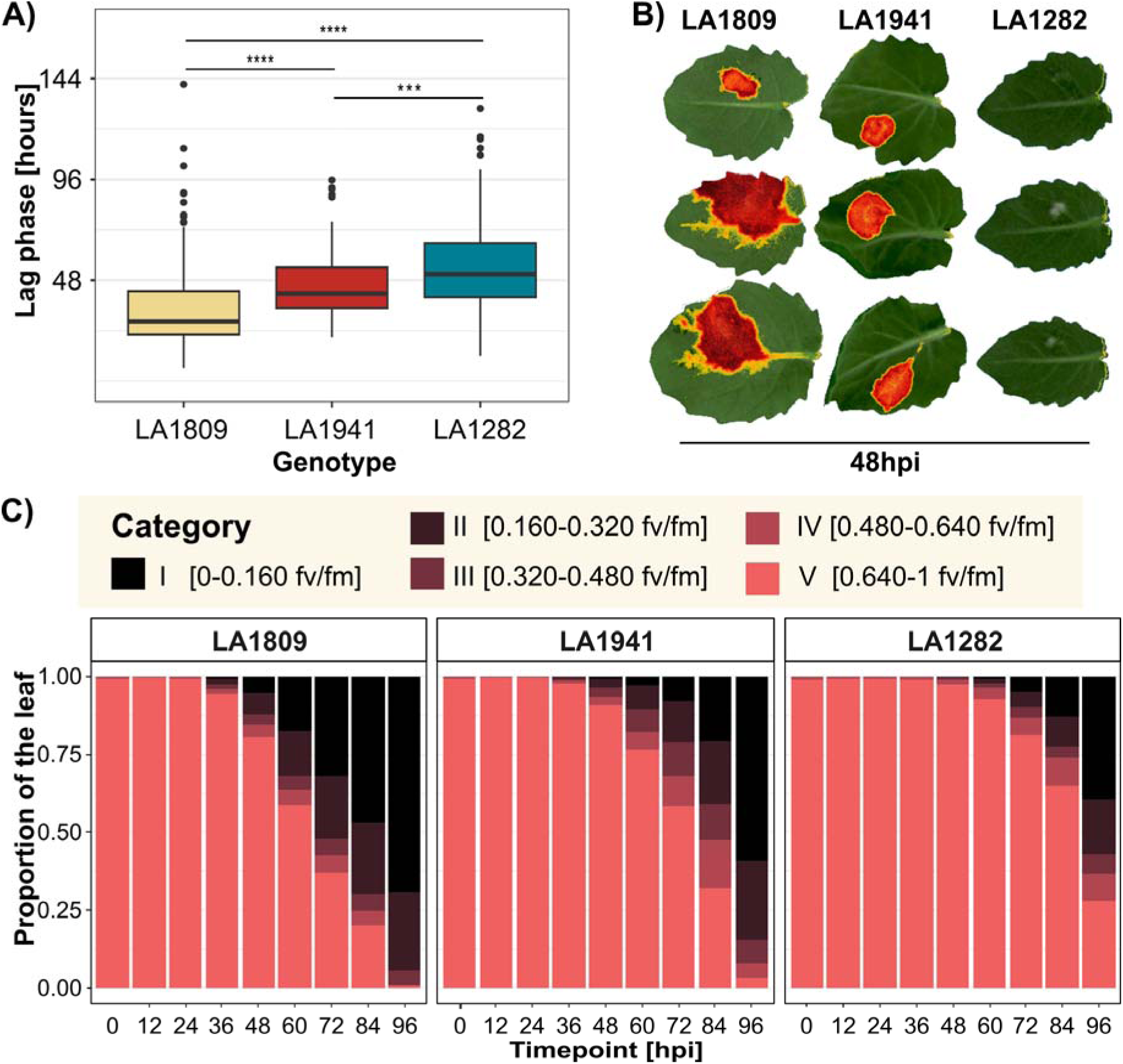
Genotypic Variation in Lag-Phase Duration and Photosynthetic Stress Response. The time until lesion onset varies significantly among the three *S. pennellii* genotypes: LA1282 (n = 205), LA1809 (n = 148), and LA1941 (n = 98). We previously assessed lag-phase duration in a screening study of *Solanum* accessions and now present the duration for these three focal genotypes. The y-axis indicates the asymptomatic phase (in hours after infection). Stars denote statistical significance from pairwise Wilcoxon tests: ***p ≤ 0.001, **p ≤ 0.0001 (**A**, data adapted from Einspanier et al. 2024). We reproduced the lag-phase phenotype using the “PlantExplorer Pro” phenotyping platform, measuring dark-adapted chlorophyll fluorescence (Fv/Fm) every 12 hours. **(B)** A representative 48 hpi overlay visualises photosynthetic stress areas, where non-stressed regions appear in true-color RGB, while regions with reduced Fv/Fm are highlighted in yellow-to-red overlay for illustration. **(C)** Stacked bar plots show the mean categorical distribution of Fv/Fm values in a time series experiment across the three genotypes. We performed two independent experiments, each with 18 leaflets per genotype and treatment.

To validate these findings and deepen their biological significance, we assessed stress-induced changes in photosynthetic efficiency by measuring the ratio of variable to maximum chlorophyll *a* fluorescence (Fv/Fm). We hypothesised that early photosynthetic stress could indicate the plant’s defence capacity and, in turn, affect the duration of the lag phase. To test this, we conducted a phenotyping experiment using detached leaves, monitoring Fv/Fm at 12- hour intervals to capture dynamic changes in photosynthetic efficiency over time. The most pronounced differences in photosynthetic stress among the genotypes emerged 48 hours post-inoculation (hpi). In LA1282, Fv/Fm remained largely unaffected, whereas LA1809 and LA1941 exhibited distinct regions of reduced fluorescence (fig. 1B). Interestingly, we observed a halo with intermediate Fv/Fm values (0.48 - 0.64) surrounding areas of high photosynthetic stress (Fv/Fm < 0.48). This suggests a narrow intersection of intermediate photosynthetic stress between healthy and diseased tissues, marked by slightly reduced photosynthetic stress levels. We further assessed the proportions of Fv/Fm categories per genotype to quantify the extent of photosynthetic stress, independent of leaf size effects. While we observed genotype-dependent temporal dynamics of the Fv/Fm classes, we found the first changes in the Fv/Fm approximately at the same time as the lesion appeared. In LA1809 first stress was measured at 36 hpi (lsmean lag phase: 38.16 h). We observed the first drops in Fv/Fm on LA1941 at 40hpi (lag phase duration 49 h), while photosynthetic stress appeared the latest in LA1282 at 60 hpi (lag phase duration 56.88h). Accordingly, we conclude that *S. sclerotiorum* inoculation leads to photosynthetic stress, which is mostly associated with lesion formation. Consequently, Fv/Fm reflects the visual phenotypes if measured on 12 h rhythms (see fig. 1C).

We observed that the proportion of transient Fv/Fm categories (i.e., categories II, III, or IV) remains stable throughout the infection process across all genotypes. Notably, the intermediate stress zone (class IV) was relatively narrow, with a transition to severe stress and eventual total loss of chlorophyll fluorescence. Over time, the ‘dead’ class (class I, Fv/Fm < 0.160) progressively expanded, while transient classes remained localised at the advancing lesion edge. Contrary to our expectations, we did not measure photosynthetic stress on infected yet asymptomatic areas of the leaf. This observation is persistent even during the lesion growth phase, suggesting that the photosynthetic stress response to *S. sclerotiorum* inoculation is spatially restricted to regions of direct pathogen interaction. Other QDR phenotypes, such as LDT, might be influenced by regulatory variability in this intersection (Sucher et al. 2020).

### Transcriptomic response to infection underlies genotype-dependent longitudinal dynamics

We performed an RNA sequencing experiment to investigate how longitudinal infection-induced gene expression dynamics influence the lag phase duration. First, we mapped 3’UTR sequencing reads to the *S. pennellii* reference genome, resulting in consistently high mapping rates in mock samples similar to other studies (85% - 97.81%, fig. 2A, Sucher et al. (2020)). The mapping rates in infected conditions gradually declined over time (70% - 97% fig. 2A), likely due to the increasing number of fungal reads reflecting the pathogen proliferation during pathogenesis (see suppl. fig 1 for mapping rates on the *S. sclerotiorum* genome). We conducted a principal components analysis (PCA) on the mapped reads to assess the overall data structure concerning plant genotype and inoculum (fig. 2). In the PC1 vs PC2 projection (fig.2B), we observed a clear separation between later-stage infected samples and the other samples, indicating a strong infection-driven effect. Although more distinct based on their lag-phase derived resistance, LA1282 and LA1809 clustered more closely together, suggesting a higher degree of transcriptome similarity between these genotypes. In the PC2 vs PC3 space (fig. 2C), the three genotypes were more distinctly differentiated, appearing approximately equidistant. These patterns indicate that the dataset captures biologically meaningful variation, with PC1 distinguishing time-dependent inoculation effects and PC2&3 genotype-specific transcriptomic differences.

**Figure 2:**
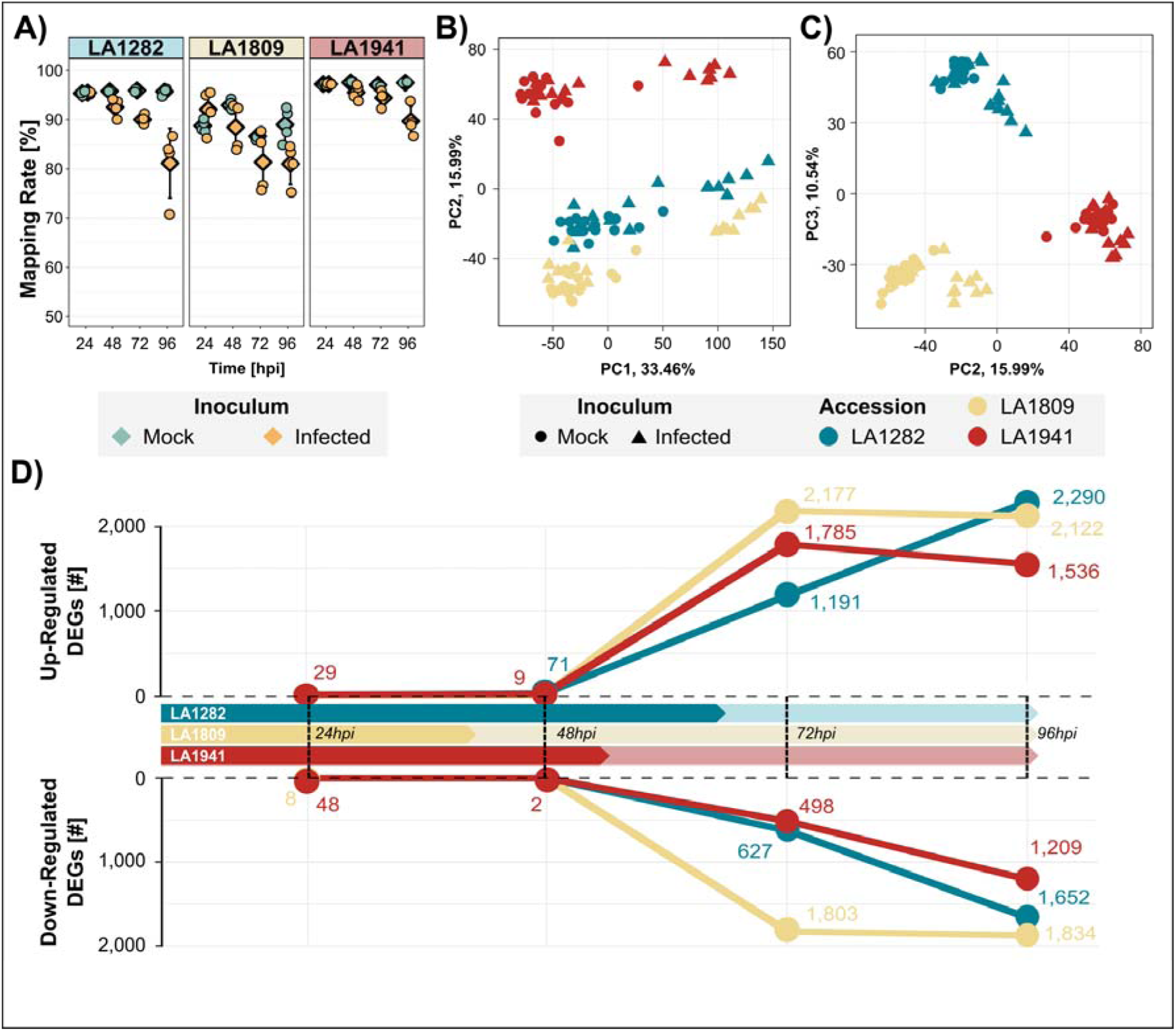
*S. sclerotiorum* infection induces gene expression shifts in all *S. pennellii* genotypes. **(A)** Mapping rates of all samples to the *S. pennellii* reference genome. The gradual decline in inoculated samples over time reflects the increasing number of fungal reads during infection. The whiskers represent the standard deviation, while diamonds indicate the mean. **(B, C)** Principal Component Analyses (PC1 vs. PC2 and PC2 vs. PC3) show clear treatment-dependent expression shifts over time **(B)**, and genotype separation **(C). (D)** Differential gene expression analysis (infection vs. mock, at each time point) reveals distinct longitudinal regulation patterns. The x-axis corresponds to the respective time points (24-, 48-, 72-, and 96-hours past inoculation), and the colour indicates the genotype. The y-axis defines the number of differentially expressed genes (|log2FC| > 1 and p-adj < 0.05) of up- (top panel) and downregulated genes (bottom panel). The solid arrow indicates the approx. mean lag phase duration of each genotype. If no dot/number is shown, no DEGs were found for that genotype at that particular time point.

We then performed a differential gene expression analysis, extracting the number of differentially expressed genes upon fungal infection (DEGs, defined as p-adj. < 0.05, |log2FoldChange| > 1) on each time point for all genotypes. Overall, we observed that the number of DEGs (contrast infection vs mock) increased for all genotypes over time. At 24 hpi, LA1941 showed 29 upregulated and 48 downregulated genes, while LA1809 exhibited only 8 downregulated DEGs. Surprisingly, we found no DEGs at 24 hpi on the most resistant genotype LA1282. By 48hpi we measured 71 upregulated DEGs on the genotype LA1282, while we found 11 DEGs on LA1941 (9 up-, 2 downregulated). At this time point, we did not detect any DEGs on the most susceptible genotype LA1809. It is unclear if the absence of detected DEGs during early infection results from host-pathogen interactions being masked by regulatory processes in uninfected tissues (fig. 2D).

We observed a stronger regulatory response once the lag phase passed, during the progression of lesion formation. Accordingly, all genotypes displayed a considerable number of DEGs at 72 hpi (LA1282 1,191 up- and 627 downregulated, LA1809 2,177 up- and 1,803 downregulated, and LA1941 1,785 up- and 498 downregulated) and 96 hpi (LA1282 2,290 up- and 1,652 downregulated, LA1809 2,122 up- and 1,834 downregulated, and LA1941 1,536 up- and 1,209 downregulated). However, we could not establish a connection between the late regulatory response (72 hpi and later) and the level of lag-phase-mediated QDR. Therefore, we hypothesise that late regulatory adjustments triggered by infection may impact other QDR phenotypes, such as the lesion growth rate or result from plant tissue degradation. Still, we observed a weak genotype-dependent differential response in early time points after *S. sclerotiorum* inoculation, with the most pronounced regulatory shift in LA1282 shortly before the lag phase ends.

### LA1282 shows significant transcriptomic reprogramming before lesion onset

Next, we focused on the 48 hpi time point, hypothesising that regulatory shifts during the lag phase could determine its duration. Notably, inoculated LA1282 leaves remain asymptomatic at this stage, whereas LA1809 leaves already show visible symptoms (fig. 1).

In total, we found 71 differentially expressed genes in LA1282 (fig. 2D). Among these DEGs we found mostly defence-associated genes, including several pathogenesis-related (PR) genes such as PR-4 (two paralogs: Sopen01g040940, log2FC=1.5, and Sopen01g040950, log2FC=1.17), PR-1 protein 38 (two paralogs: Sopen01g026170, log2FC=1.43, and Sopen01g048970, log2FC=1.21) and the strongly infection-induced protein LOX1 (log2FC=4.3). We also identified proteins involved in the protection against oxidative stress (DOX1, log2FC=3.95), interaction with fungal pathogens (e.g., chitinases, and β-1,3- endoglucanases), and detoxification (glycosyltransferases, cytochrome p450). Among those DEGs were several transcription factors, such as WRKY45 and WRKY51, as well as ethylene-response factors, illustrating a broad regulatory response to the inoculum. In addition to those well-characterized defence genes, we found several proteins of unknown function, including Sopen08g008720, Sopen12g023430, and Sopen03g007540. Interestingly, we did not detect any genes down-regulated in this genotype at 48 hpi (suppl. tab. 1, fig. 3A).

**Figure 3:**
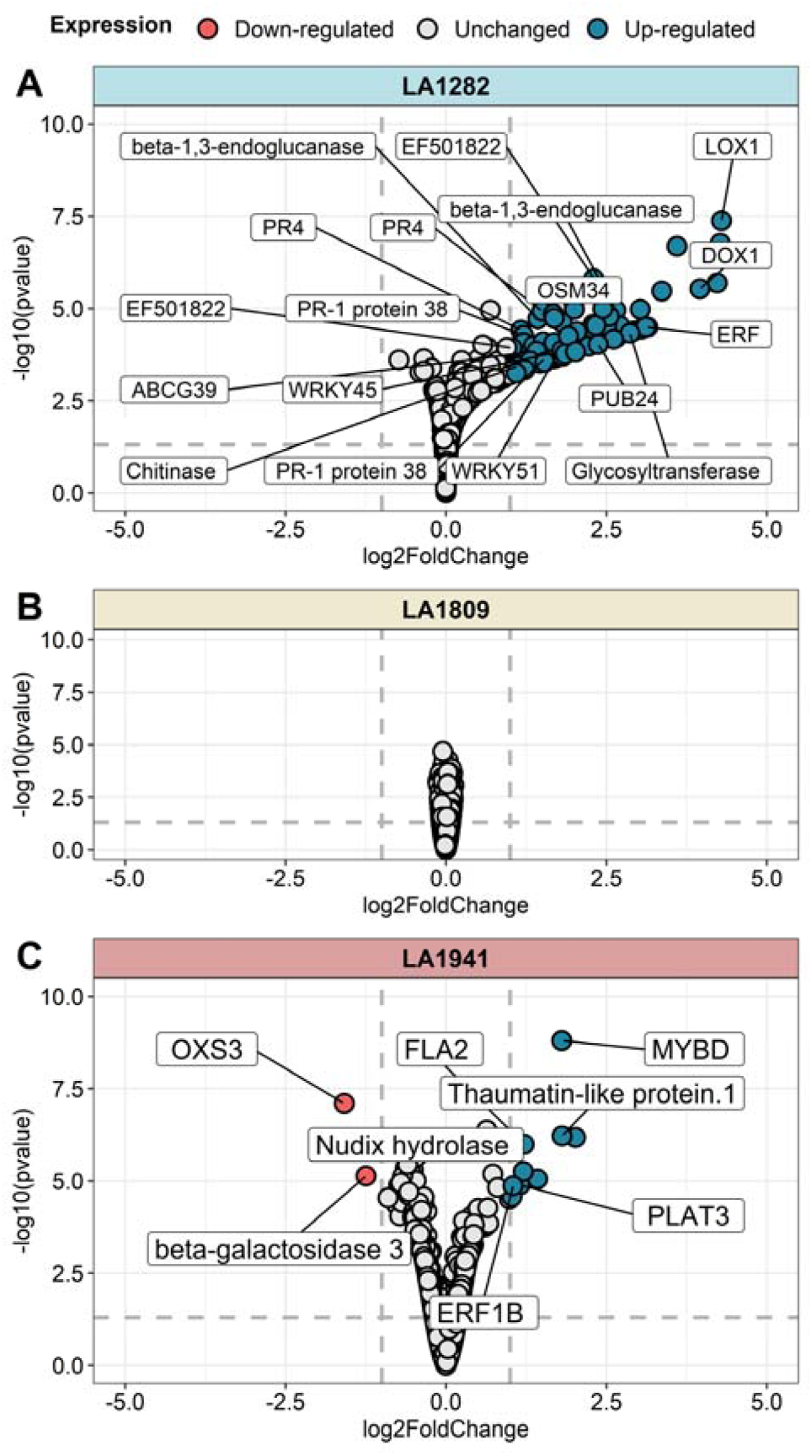
LA1282 exhibits distinct regulatory reprogramming at 48 hpi. The volcano plots display differential gene expression analysis results comparing infected vs mock conditions at 48 hpi. We tested the genotypes LA1282 **(A)**, LA1809 **(B)**, and LA1941 **(C).** Each dot represents a single gene. The y-axis shows the - log_10_ of the FDR-corrected p-value, while the x-axis represents the log_2_ fold-change. The dotted lines mark the thresholds for DEG assignment (|log_2_ fold-change| > 1 and p-adj. < 0.05). Genes meeting these criteria are considered differentially expressed. Red dots indicate downregulated DEGs, while blue dots highlight genes with increased expression in infected conditions. Labelled genes indicated transcripts of specific interest.

Although the average lag phase duration of LA1809 ends prior to 48hpi, we did not detect any statistically significant regulatory reprogramming. This finding suggests an incomplete, or delayed response that lags behind the infection process (fig. 2D, fig. 3B).

In contrast, we identified two downregulated and nine upregulated DEGs at 48 hpi in the intermediate-resistant genotype LA1941. Among these, we detected MYB (log2FC=1.8), and ERF (log2FC=1.05) transcription factors as well as genes directly linked to pathogenesis. These include a putative lipoxygenase (PLAT3, log2FC=1.15) and a putative PR-5 family protein (thaumatin-like protein 1, log2FC=1.8). Interestingly, we found that OXIDATIVE STRESS 3, a key regulator of ROS homeostasis and stress resilience, was significantly downregulated (log2FC = -1.5) at this time point (suppl. tab. 2, Fig. 3C).

### Gene Set Enrichment Analysis on LA1282 at 48hpi reveals clear defence responses

To further validate the early transcriptomic reprogramming in the resistant genotypes, we performed a gene set enrichment analysis (GSEA). This approach allows for a functional interpretation of subtle regulatory changes by analysing ranked gene expression patterns across the entire transcriptome rather than focusing solely on DEGs.

We observed that the transcriptome of the susceptible genotype LA1809 is largely dominated by basal homeostasis processes at 48hpi. This includes the processes “translation” (GO:0006412, NES=4.05), “ribosomal small subunit assembly” (GO:0000028, NES=3.79), and “energy reserve metabolic processes” (GO:0006112, NES=2.88, suppl. tab. 3). This pattern is further reinforced by the absence of defence-related processes in the LA1809 GSEA analysis 24hpi. These insights further highlight how LA1809 cannot mount a robust defence response both before and after lesion onset (suppl. tab. 3). .

In accordance with our previous findings, we observed a significant enrichment of what has been defined as defence-related processes in the most resistant genotype, LA1282, at 48hpi. This set includes “response to fungus” (GO:0009620, NES=2.9), “ethylene metabolic process” (GO:0009692, NES=2.79), and the jasmonic acid-associated “oxylipin biosynthetic process” (GO:0031408, NES=2.83). We also identified a more general “defence response” (GO:0006952, NES=2.09, set size=151). We found evidence for secondary defence-related processes, such as “lipid biosynthetic processes” (GO:0008610, NES=2.85) and “carbohydrate metabolic processes” (GO:0005975, NES=2.38), which might reflect the plant’s ability to recognise and metabolise fungal structures such as chitin or glucans. Interestingly, we detected no defence-related enriched gene sets at 24 hpi. Although these enrichments largely mirror the DEGs observed in this genotype, we argue that the significantly expanded defence-associated gene set indicates a broad transcriptomic reprogramming, involving a larger number of genes under nuanced regulatory control (tab.1).

**Table 1.**
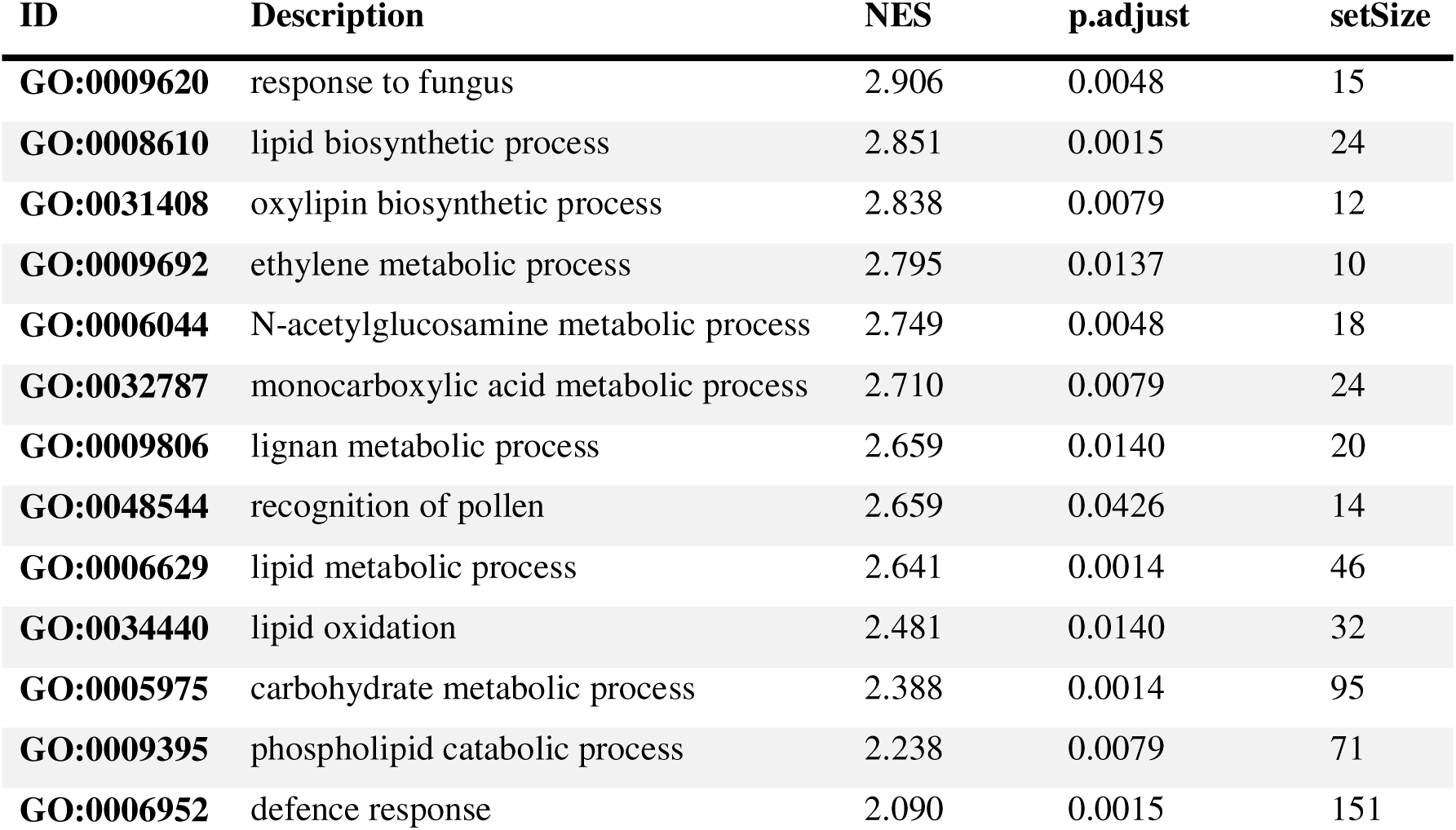
Gene Set Enrichment Analysis (GSEA) of ranked log2FC gene expression data in LA1282 at 48hpi. All genes were ranked according to their differential expression, and significant enrichments (p-adj. < 0.05) GO terms (biological process) were identified. The normalised enrichment score (NES) measures the magnitude of enrichment of a given gene set relative to its size and compares enrichment across different gene sets. P-values were FDR-corrected using the Benjamini-Hochberg procedure.

| **ID** | **Description** | **NES** | **p.adjust** | **setSize** |
| --- | --- | --- | --- | --- |
| **GO:0009620** | response to fungus | 2.906 | 0.0048 | 15 |
| **GO:0008610** | lipid biosynthetic process | 2.851 | 0.0015 | 24 |
| **GO:0031408** | oxylipin biosynthetic process | 2.838 | 0.0079 | 12 |
| **GO:0009692** | ethylene metabolic process | 2.795 | 0.0137 | 10 |
| **GO:0006044** | N-acetylglucosamine metabolic process | 2.749 | 0.0048 | 18 |
| **GO:0032787** | monocarboxylic acid metabolic process | 2.710 | 0.0079 | 24 |
| **GO:0009806** | lignan metabolic process | 2.659 | 0.0140 | 20 |
| **GO:0048544** | recognition of pollen | 2.659 | 0.0426 | 14 |
| **GO:0006629** | lipid metabolic process | 2.641 | 0.0014 | 46 |
| **GO:0034440** | lipid oxidation | 2.481 | 0.0140 | 32 |
| **GO:0005975** | carbohydrate metabolic process | 2.388 | 0.0014 | 95 |
| **GO:0009395** | phospholipid catabolic process | 2.238 | 0.0079 | 71 |
| **GO:0006952** | defence response | 2.090 | 0.0015 | 151 |

At 24 hpi, we observed a significant enrichment of “response to fungus” (GO:0009620, NES = 3.05, set size = 11) and “cellular detoxification of aldehyde” (GO:0110095, NES = 2.87) in the LA1941 transcriptome (tab. 2). However, by 48 hpi, only three processes remained significantly enriched: “defence response” (GO:0006952, NES = 2.92), “cytoplasmic translation” (GO:0002181, NES = 3.24), and “microtubule-based movement” (GO:0007018, NES = 3.1; suppl. tab. 4). These findings suggest an early activation of pathogen recognition and detoxification mechanisms, followed by a later shift toward defence activation, protein synthesis, and cytoskeletal rearrangements.

**Table 2.** Gene Set Enrichment Analysis (GSEA) of ranked log2FC gene expression data in LA1941 at 24hpi. All genes were ranked according to their differential expression, and significant enrichments (p-adj. < 0.05) GO terms (biological process) were identified. The normalised enrichment score (NES) measures the magnitude of enrichment of a given gene set relative to its size and compares enrichment across different gene sets. P-values were FDR-corrected using the Benjamini-Hochberg procedure.

| ID | Description | NES | p.adjust | setSize |
| --- | --- | --- | --- | --- |
| GO:0006457 | protein folding | 3.388 | 0.00014 | 82 |
| GO:0006412 | translation | 3.095 | 0.00014 | 98 |
| GO:0009620 | response to fungus | 3.055 | 0.02500 | 11 |
| GO:0006730 | one-carbon metabolic process | 3.005 | 0.03705 | 10 |
| GO:0009813 | flavonoid biosynthetic process | 2.900 | 0.02500 | 23 |
| GO:0110095 | cellular detoxification of aldehyde | 2.879 | 0.00563 | 54 |
| GO:0006516 | glycoprotein catabolic process | 2.579 | 0.03705 | 51 |
| GO:0000028 | ribosomal small subunit assembly | 2.311 | 0.01930 | 121 |

These findings suggest that the high-QDR genotypes LA1282 and LA1941 exhibit infection-induced regulatory changes towards defence processes. Notably, there is little overlap between the two genotypes: LA1941 initiates a subtle yet earlier response characterised by a small set size and weakly differentially expressed genes. In contrast, LA1282 initiates a later but more robust defence response, with a larger set size and a greater number of strongly regulated DEGs, particularly at 48 hpi. The lack of overlap between these responses suggests that each genotype employs independent regulatory mechanisms to achieve quantitative disease resistance (QDR). Meanwhile, LA1809 fails to mount any detectable response during early pathogenesis, further underscoring its susceptibility.

### The resistant accession LA1282 shows distinct regulation of putative resistance-associated genes

We hypothesised that the early induction of pathogenesis-related genes might confer a QDR advantage to LA1282. Based on our previous findings, we selected the 71 DEGs identified in LA1282 at 48 hpi for further analysis, hereafter referred to as “focal DEGs” (see fig. 3A). Following this, we expect increased susceptibility associated with altered temporal orchestration, such as delayed or absent expression of the focal DEGs. To test this hypothesis, we examined the longitudinal (temporal) trajectories of the focal genes across the three genotypes.

Firstly, we performed a PCA on the per-genotype log2 fold-change (infected vs mock) across all time points for each focal DEG, revealing a clear separation among the genotypes. Each dot in the PCA represents a single focal DEG in a specific genotype, encompassing its regulation across all four time points. Notably, LA1809 and LA1941 cluster more closely, with some trajectories overlapping, whereas LA1282 trajectories were more separated (fig. 4A).

**Figure 4:**
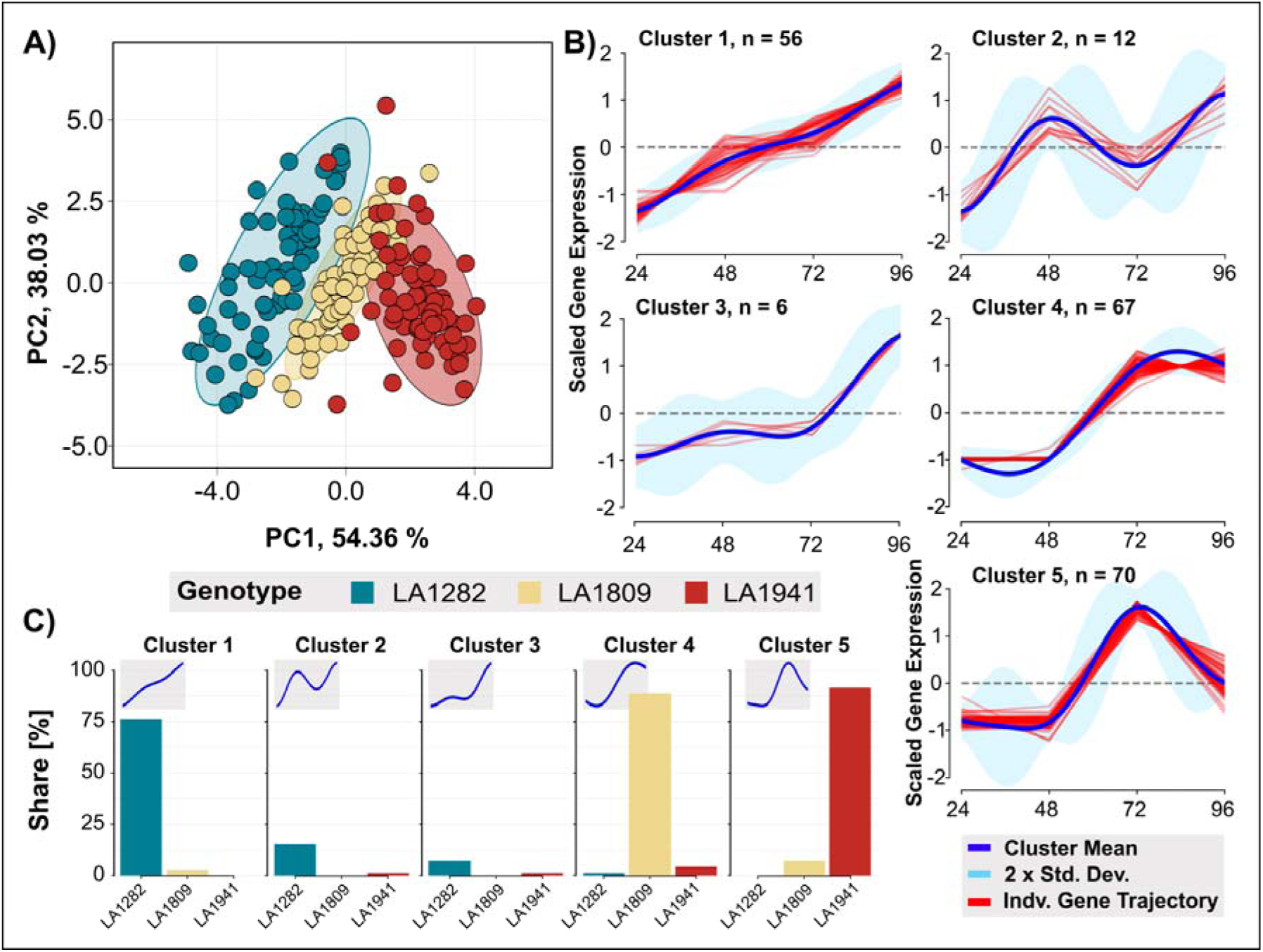
LA1282-specific early response DEGs underlie divergent temporal trajectories in more susceptible genotypes. **A)** Principal component analysis (PCA) of the 71 focal gene trajectories across the three genotypes. Each dot represents the temporal trajectory of a single focal DEG (incorporating the four time points). Colours represent the genotype identity, while the ellipses denote each genotype’s 95 % confidence interval. **B)** Clustering analysis of the genotype-dependent trajectories of the 71 focal genes. A Dirichlet Process Gaussian Process model was used to determine the ideal number of clusters and model the individual gene trajectories over time. The algorithm identified five clusters. The y-axis defines the scaled gene expression, while the x-axis denotes the time. The blue line represents the mean trajectory of each cluster, while the red lines show individual gene trajectories. The shaded regions represent two standard deviations (2x SD). The number of genes assigned to each cluster is indicated by n. **C)** Cluster membership analysis of the focal gene trajectories per genotype. The y-axis represents the per-genotype proportion of focal DEGs assigned to each cluster. The colour and x-axis define the different genotype clusters, corresponding to those defined in (B).

We performed a temporal clustering analysis to describe the regulatory dynamics among the three genotypes. For this, we analysed the longitudinal expression pattern of the 71 focal DEGs per genotype (3 x 71 focal DEGs). All focal DEGs show clear evidence of expression in each genotype at least once during our time-series experiment. Using a Dirichlet Process Gaussian Process Model (DP_GP), we grouped trajectories into clusters based on their shared temporal expression profiles, allowing us to determine whether the same genes followed similar or divergent regulatory patterns across genotypes.

Through this analysis, we identified five distinct expression patterns (fig. 4B). Cluster 1 includes 56 genes with a continuous increase in expression from 24hpi to 96hpi. In contrast, the trajectory of the 12 genes in cluster 2 followed a zigzag pattern with expression peaks at 48hpi and 96hpi. Cluster 3, containing only six genes, shows a modest rise at 48hpi followed by a stronger increase till 96hpi. The remaining gene trajectories fell into Cluster 4 (n=67) and Cluster 5 (n=70). Both clusters show increased expression after 48 hpi, but Cluster 5 peaks and drops sharply at 72hpi, while Cluster 4 plateaus at a maximum.

Consequently, we grouped the gene trajectories of the focal genes into three major regulatory categories:

1. Early-response genes (clusters 1–3)
2. Late-expression trajectories with sustained expression (cluster 4)
3. Late-expression gene trajectories with transient expression peaking at 72hpi before declining (cluster 5).

We performed a membership analysis of cluster assignment to test whether LA1282 trajectories exhibit an early increase, while those of LA1941 and LA1809 are expressed late. Strikingly, all but one of the LA1282 trajectories clustered within the early response group (Clusters 1-3). In contrast, most focal DEG trajectories of LA1809 and LA1941 fell into the late-responding Clusters 4 and 5.

Interestingly, most LA1941 trajectories showed clear downregulation at 96 hpi, whereas those genes remained highly upregulated in LA1809. We observed very little overlap between the genotypes, strengthening the hypothesis that the longitudinal expression dynamics are highly genotype-specific (fig. 4C). Although LA1282 trajectories were distributed across multiple clusters, LA1809 genes predominantly grouped in Cluster 4, while LA1941 genes primarily fell into Cluster 5, again suggesting distinct regulatory strategies.

To summarise, all focal DEGs show infection-induced regulation in all genotypes, highlighting that LA1809 susceptibility is not due to the absence of the focal DEGs from its regulatory repertoire. Moreover, we observed that the focal DEGs underlie clear genotype-dependent and infection-induced temporal dynamics in the tested genotypes. Building on these results, we hypothesise that ulatory trajectory of focal DEGs (zig-zag pattern), characterised by subtle early induction followed by (partial) deregulation.

### Basal expression of a WRKY transcription factor might determine the level of QDR

Building on the high genotypic specificity of focal DEG trajectories, we hypothesised that these genes might be regulated by a shared transcription factor (TF). We performed a directed gene regulatory network analysis to test this, identifying 28 potential regulatory hub genes.

We identified one transcription factor (Sopen02g011350) with regulatory connections to all 71 focal genes. Notably, this TF is not part of the focal DEGs.

Interestingly, Sopen02g011350 is an ortholog of the *A. thaliana* WRKY6, a TF with a documented role in resistance, primarily against abiotic stresses (Chen et al. 2009; Huang et al. 2016; Niu et al. 2024). We found that Sopen02g011350 had a significantly elevated eigencentrality in the GRN, indicating a key role in network regulation. Additionally, this TF displayed a high edge weight to all focal DEGs, supporting a potential function as a central regulator of the focal DEGs.

Next, we analysed the expression pattern of this TF in both mock and infected conditions across three genotypes to verify its role in genotype-dependent longitudinal regulation. Compared to LA1809 and LA1941, LA1282 exhibited consistently higher expression of Sopen02g011350, a trend observed in both infected and non-infected conditions, independent of time. Notably, under mock conditions at 24 hpi, LA1282 expressed Sopen02g011350 significantly higher than the other genotypes. Upon infection, Sopen02g011350 expression increased in all genotypes, though its basal expression level in LA1282 remained consistently elevated (fig. 5, suppl. table 9).

**Figure 5:**
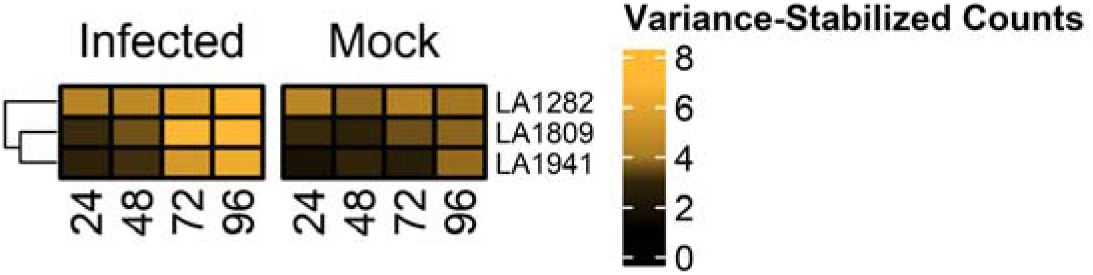
The gene regulatory network hub Sopen02g011350 (WRKY6 ortholog), exhibits high basal and induced expression in the resistant genotype LA1282. We identified Sopen02g011350, a putative WRKY6 ortholog, as a gene regulatory network (GRN) hub based on high eigencentrality and connectivity to all focal DEGs. We analysed its expression using mean variance-stabilised of absolute (deduplicated) read counts. The heatmap displays Sopen02g011350 expression levels, where yellow indicates high expression and black represents low expression. We compared four time points (24–96 hpi) under infected and mock conditions across the three genotypes: LA1282, LA1809, and LA1941.

### Basal expression of focal DEGs is significantly elevated in LA1282

Following the observation that the putative core regulator of the focal DEGs (Sopen02g011350) exhibited significantly higher basal expression in LA1282, we next investigated whether the focal DEGs also show constitutively elevated expression in this genotype.

Notably, most focal DEGs (56 out of 71, suppl. tab. 9) are significantly more expressed in LA1282 mock conditions than the susceptible genotype LA1809. In particular, pathogenesis-related genes PR-1, PR-4, and β-1,3-endoglucanases displayed stronger basal expression (fig. 6, suppl. tab. 9). This further provides evidence that basal or preformed defence mechanisms contribute to QDR in LA1282.

**Figure 6:**
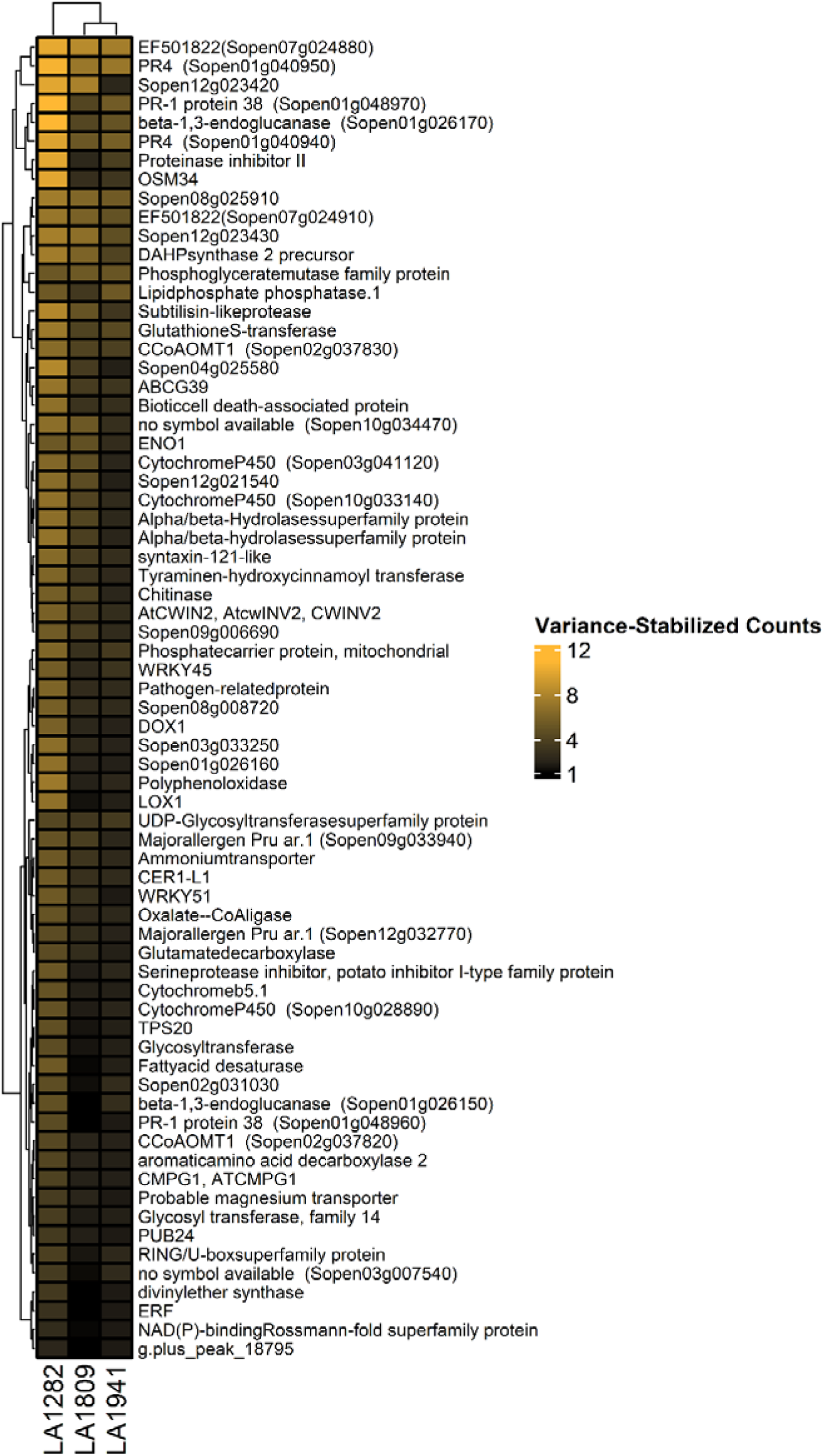
LA1282 exhibits higher basal expression of Focal DEGs. We analysed the basal expression of the 71 focal genes (identified in LA1282 at 48 hpi) by comparing their expression under mock conditions across the three genotypes. We used variance-stabilised mean expression values, which were min-max normalised to the highest and lowest observed values. The heatmap visualises gene expression levels, where yellow indicates the highest expression and black denotes the lowest. The expression values represent measurements at 24 hpi under mock conditions across four independent biological replicates. Rows and columns were clustered using Euclidean distance.

Since LA1282 exhibits elevated basal expression of both the core regulator Sopen02g011350 and its downstream focal DEGs, we next asked whether pathogen perception might contribute to QDR in this genotype. To explore this, we analysed the basal expression of key pathogen recognition proteins, including BAK1, SOBIR1, CERK1, receptor-like proteins (RLPs), and receptor-like kinases (RLKs).

Interestingly, we found eight RLKs with significantly elevated basal expression in LA1282 compared to LA1809. Moreover, we identified seven differentially expressed RLPs, including three upregulated in LA1282 and one allele each of CERK1 and BAK1 (see suppl. tables 5-8). These findings illustrate the first evidence of how the basal expression of key defence genes and the pre-activation of immune receptors might facilitate an earlier and more rapid defence response in LA1282, resulting in enhanced QDR.

## Discussion

We performed a longitudinal transcriptome analysis to identify regulatory factors driving the lag phase duration of *S. pennellii* upon *S. sclerotiorum* inoculation. Our goal was to characterise the composition and temporal dynamics of the regulatory response in high-QDR genotypes. Specifically, we tested whether QDR is driven by induced or basal expression.

### The lag phase duration is determined by common defence genes

In the high-QDR genotype LA1282, we identified a set of focal DEGs which we associated with resistance (fig. 2D). These genes show evidence for late infection-induced expression (72-96hpi) in more susceptible genotypes yet lack the longitudinal pattern found in LA1282 (fig. 4B). In particular, the genotype LA1282 facilitates early defence gene expression at 48hpi, before lesion onset. Many of these focal DEGs, including glutathione S-transferases, ABC-transporters, p450 cytochromes, receptor-like kinases/proteins and PR-genes, have been previously linked to *S. sclerotiorum*-induced defence responses across multiple hosts (Xue et al. 2024; Sucher et al. 2020; Wei et al. 2016; Gullner et al. 2018). These genes contribute to necrotrophic pathogen defence by regulating detoxification, reactive oxygen species homeostasis, transport and cell death (Xue et al. 2024).

While general defence responses to *S. sclerotiorum* inoculation are well understood, comparative studies examining multiple genotypes with varying levels of QDR remain scarce. Little is known about how temporal regulation influences QDR phenotypes. However, some reports have linked early gene expression patterns to QDR against *S. sclerotiorum* on *Brassica juncea* or soybean (Que et al. 2014; Ranjan et al. 2019; Yang et al. 2025). Ranjan et al. (2019) observed that early jasmonate (JA) accumulation enhances ROS scavenging capacity in a resistant genotype, which aligns with our findings on LA1282. The role of JA in conferring resistance to *S. sclerotiorum* has been demonstrated, although the significance of longitudinal JA regulation remains inconclusive (Wei et al. 2016). While some studies do not establish a direct link between temporal gene regulation and resistance (Seifbarghi et al. 2017) or remain inconclusive (Chang et al. 2018), our findings align with prior work indicating that focal gene activity is crucial for lag-phase-mediated resistance (Westrick et al. 2019). We observed that a distinct regulatory shift at 48hpi is required to delay lesion onset on a high- QDR genotype.

Studies on *S. sclerotiorum* point to a distinct regulatory shift between the early (presymptomatic, 24-48hpi) and late (symptomatic, 96hpi) stages of infection, potentially marking the transition from biotrophy to necrotrophy (Seifbarghi et al. 2017; Westrick et al. 2019; Kabbage et al. 2015). These presymptomatic gene-expression changes align closely with our observation that a high-QDR genotype (LA1282) initiates an early regulatory response before lesion onset, suggesting strong genotype-by-genotype interactions (Rowe and Kliebenstein 2010). This could explain the spatio-temporal variation, potentially affecting the length of the biotrophic phase, along with host responses to fungal attack strategies that shape QDR (fig. 1C & fig.2).

### A WRKY Transcription Factor might control the rapid QDR response to *S. sclerotiorum* inoculation

The role of WRKY TFs in oxalic acid and *S. sclerotiorum* resistance is well documented. Multiple WRKY TFs have been identified across multiple hosts like *A. thaliana,* oil seed rape, or *Brassica juncea*, including AtWRKY28, AtWRKY75, BnWRKY8, BnWRKY33, BnWRKY61 or the WRKY25 *BjuA016484* (Chen et al. 2013; Wei et al. 2016; Yang et al. 2025; Zhao et al. 2024; Wang et al. 2014). In this study, gene regulatory network and differential gene expression analysis revealed WRKY6 (Sopen02g011350) as a putative regulator of QDR-related genes. Interestingly, WRKY6 has been characterised in *A. thaliana* for its role in abiotic stress resistance, particularly in response to phosphorus or potassium limitation (Chen et al. 2009; Niu et al. 2024). However, some studies have reported its differential expression upon *S. sclerotiorum* infection in *A. thaliana* and oil seed rape (Zhang et al. 2022; Wu et al. 2016). We speculate that the association of WRKY6 with abscisic acid (ABA) homeostasis might enable its role in defence via plant hormone signalling (Huang et al. 2016).

In tomato (*S. lycopersicum)*, WRKY6 (Solyc02g080890) is linked to pathogen resistance, with elevated expression reported in genotypes resistant against *Ralstonia solanacearum* and *Fusarium oxysporum* f. sp. *lycopersici* (Zhang et al. 2020; López et al. 2021). In contrast, in *S. pennellii*, WRKY6 has so far only been associated with leaf thickness development (Coneva et al. 2017).

Given its role in multiple stress responses and pathogen interactions, WRKY6 may serve as a key factor in resistance to diverse pathogens, consistent with observations of the typical pleiotropic nature of QDR-loci (Roux et al. 2014). We demonstrate that basal WRKY6 regulation influences complex QDR phenotypes, such as lag-phase duration, without conferring full resistance. Since WRKY6 exhibits strong regulatory connectivity to all focal genes and shows significantly elevated basal expression in the resistant genotype, similar to the focal genes, we propose that WRKY6 is a promising candidate for further investigation within the genetic regulatory network governing QDR-associated genes.

Following the canonical model of defence response to most pathogens, TFs, like WRKYs and ERFs, act downstream of key components in the pattern-triggered immunity (PTI) signalling cascade. This includes early-response elements such as mitogen-activated protein kinases (MAPKs), as well as ROS bursts and Ca^2+^ signalling, which can contribute to QDR against *S. sclerotiorum* (Li et al. 2016; Lewis et al. 2015; Wei et al. 2016; Zhang and Klessig 2001; Lin et al. 2024; Wu et al. 2016). Unfortunately, the resolution of our time-series experiment does not capture the earliest events in the host-parasite interaction. Nevertheless, we observed clear differential regulation of several potentially PTI-associated genes in the resistant genotypes, supporting their role in QDR-associated transcriptional reprogramming.

### QDR is determined by basal gene expression and defence priming

Based on our observations, we propose that elevated basal expression of focal genes, possibly through a primed defence state, and altered pathogen perception through receptor-like proteins and -kinases determine QDR in conjunction. Consistent with this hypothesis, some of our candidate genes exhibit increased basal expression in resistant oilseed rape genotypes, resulting in stronger induction compared to susceptible backgrounds (Wu et al. 2016). We found significant variation in the basal RLK and RLP regulation across genotypes with varying QDR levels. Nonetheless, these differences in RLK and RLP trajectories require a more detailed pathway analysis and molecular characterisation assays, such as ROS burst or phytohormone measurements, to gain insight into the pathways regulating the link between basal RLK/RLP expression and the regulation of the focal DEGs.

Notably, the high-QDR genotype LA1282 shows a strong basal expression of the receptor-like kinase LIK1 (Sopen02g020810). In *A. thaliana,* LIK1 is phosphorylated by CERK1 and has been shown to enhance resistance against *S. sclerotiorum* while acting as a susceptibility factor for biotrophic pathogens (Le et al. 2014). LIK1 plays a role in chitin perception, potentially interacting with FLS2, PEPR1, and PEPR2. Interestingly, LIK1 mutants show increased sensitivity to chitin (Le et al. 2014), but fail to respond to glycans, suggesting that LIK1 mediates the recognition of carbohydrate-based elicitors, particularly mixed-link glucans (Martín-Dacal et al. 2023). Beyond RLKs, RLPs also contribute to QDR, as demonstrated by AtRLP30, AtRLP42 and AtRLP23, which mediate fungal elicitor perception (e.g., SCFE1 or nlp20), increasing QDR against multiple pathogens (Mbengue et al. 2016; Albert et al. 2015). Yet, most of the receptors identified here remain uncharacterised and appear unique to wild tomatoes. Based on our results, we raise the question of whether the QDR phenotypes reported in most receptor-based studies are primarily attributed to delayed lesion onset.

## Conclusion

In this study, we observed that subtle shifts in gene expression can significantly extend the lag-phase duration. We speculate that the fungus may be particularly vulnerable during the early stages of infection, making basal expression and fine-tuned longitudinal gene expression changes essential components of QDR.

Based on the elevated abundance of focal DEGs and PRRs, we hypothesise that LA1282 might benefit from a primed state of physiological awareness. Previously, it was shown that AtWRKY6 and PR1 are subject to trans-generational defence priming, linking transgenerational systemic acquired resistance with histone modification (Conrath et al. 2015; Luna et al. 2012). The constitutively elevated defence gene expression during the primed state might enhance QDR with a relatively low fitness cost (Ahmad et al. 2010; Li et al. 2016; Conrath et al. 2006). This study demonstrates that basal gene regulation can shape QDR by modulating the temporal dynamics of a key set of defence genes, ultimately enhancing the plant’s ability to delay pathogen invasion.

## Methods

### Plant growth conditions

To characterise the regulation of lag phase duration, we selected three *S. pennellii* accessions (LA1282, LA1809, LA1941) with contrasting lag-phase duration based on findings of an earlier study (Einspanier et al. 2024). We initially acquired seed material from the C. M. Rick tomato Genetics Resource Center of the University of California, Davis (TGRC UC-Davis, https://tgrc.ucdavis.edu/). TRCG accessions can show genomic variation, as they represent a wild population and are not inbred line. Therefore, we selected one plant per accession to represent the accession genotype. After the germination, each plant was grown in the greenhouse facility of the Department of Phytopathology and Crop Protection, Faculty of Agricultural and Nutritional Sciences of the Christian Albrechts University Kiel, Germany. We maintained and multiplied adult plants via cuttings (Chryzotop Grün 0.25 %) on Stender C700 substrate to allow repeated experiments. Plants were cultivated in a controlled environment with a temperature of 21°C ± 10°C, 65% relative humidity, and a 16-hour light cycle. We applied 1% Sagaphos Blue to fertilise the plants monthly using the drip irrigation system. For the experiment, we grew 20 plants per accession/genotype from cuttings and harvested the first fully matured leaves.

### Fungal growth conditions

For our infection experiments, we utilised the generalist plant pathogen *Sclerotinia sclerotiorum* isolate 1980. We freshly cultivated the fungus from sclerotia on potato dextrose agar (PDA) at 25°C in dark conditions. To create the inoculum, we incubated four 1 cm diameter agar plugs in 100 mL of potato dextrose broth (PDB). After incubating for four days on a rotating shaker at 24°C, we mixed the suspension and vacuum-filtered it through cheesecloth. We then concentrated the resulting filtrate with empty PDB to achieve an optical density (OD) at 600 nm of OD_600_=1, using empty PDB as a mock treatment. Lastly, Tween80 was added as a surfactant to enhance the dispersion of the fungal material.

### Lag-phase duration measurement

The lag-phase duration estimates for the respective *S. pennellii* accessions were reported earlier. In short, camera-monitored detached-leaf assays were carried out in a custom-built phenotyping system called navautron (Einspanier et al. 2024; Barbacci et al. 2020). Following thresholding-based image analysis, the per-leaf lesion progression was subjected to segmented regression analysis to determine the inflexion point that marks the start of lesion growth. The time until this point defines the lag-phase duration. We repeated the experiment using multispectral imaging with the PlantExplorer Pro phenotyping platform (PhenoVation, The Netherlands) to validate our findings further. The experimental setup mirrored previous studies, except that leaves were placed on custom 3D-printed trays to facilitate transfer from Navautron conditions to the PE chamber. A 12-hour time series was conducted, and the experiment was repeated twice independently, using 18 leaves per genotype and condition. Each repetition was based on an independent plant stock.

### Measurement of Fv/Fm

We used the PlantExplorer Pro platform to measure the maximum photochemical quantum yield of photosystem II. The device was equipped with a 16MP sensor. The measure is calculated as the ratio of the variable to the maximum chlorophyll *a* fluorescence (Murchie and Lawson 2013; Xia et al. 2023). This was facilitated following a 20 min dark-adaptation of the leaves. Categorical per-leaf Fv/Fm was later determined using the PhenoVation DataAnalysis software v5.8.4-64b.

### Sequencing experiment

To capture the transcriptomic profile over the gradient of lag-phase duration, we conducted an independent navautron-like infection experiment on detached leaves. Samples were collected at 24-hour intervals (24–96 hours post-inoculation), pooling two leaves per biological replicate. We obtained four independent replicates for each time point and genotype, which were then sequenced.

We transferred the leaf samples directly into a lysis-tube containing 750 µL RNA/DNA-shield (ZymoResearch, Germany). Then, full RNA isolation was performed using the Quick-RNA Plant Miniprep kit (ZymoResearch). We measured RNA integrity and quality using gel-electrophoresis and a nanodrop one (ThermoFisher); RNA quantity was measured using a fluorometer (Promega Quantus) and a 5300 Fragment Analyzer System (Agilent). Lexogen NGS Services (Vienna, Austria) performed the library preparation and sequencing using the QuantSeq Fwd V2 UID library kit with additional unique molecular identifiers (UMIs).

### Bioinformatic processing

We assessed raw-read quality using MultiQC/FastQC, then processed the reads using the RNAseq-pipeline v3.17.0 from the Nextflow-based nf-core v24.04.2 framework. In short, reads are checked for sequencing quality and unique molecular identifiers (UMIs) extracted using UMI-tools v1.1.5. Then, adapter sequences and low-quality reads were removed with cutadapt v4.9 and trimgalore v0.6.10. rRNA depletion was skipped as no overrepresented sequences were detected. Next, the trimmed reads were aligned against the reference genome using STAR v2.7.11b followed by SALMON v1.10.3 quantification. Reads were then deduplicated using UMI-tools. We did not assemble isoforms; only gene-level expression was quantified. To accommodate 3’UTR sequencings, we executed the pipeline with custom settings:

--with_umi --umitools_extract_method "regex" --umitools_bc_pattern "^(?P<umi_1>.{6})(?P<discard_1>.{4}).*", --extra_star_align_args "-- alignIntronMax 1000000 --alignIntronMin 20 --alignMatesGapMax 1000000 -- alignSJoverhangMin 8 --outFilterMismatchNmax 999 --outFilterMultimapNmax 20 --outFilterType BySJout --outFilterMismatchNoverLmax 0.1 --clip3pAdapterSeq AAAAAAAA"

We mapped the reads against the *S. pennellii* reference genome (GenBank assembly GCA_001406875.2, updated version by Bolger et al. (2014)) and used a curated genome annotation (see online resources, Einspanier et al. 2025). During the same pipeline, we concatenated the *S. sclerotiorum* 1980 reference genome to the *S. pennellii* reference, allowing us to measure fungal gene expression (GenBank assembly GCA_001857865.1, Derbyshire et al. 2017).

We listed detailed execution reports, QC metrics and software versions in the digital resources.

### Differential Gene Expression Analysis

We used the R-package DESeq2 v1.46.0 to identify differential gene expression between the treatments or genotypes (Love et al. 2014). For the treatment effects, we converted each genotype’s count matrix into a DESeqDataSet using the `DESeqDataSetFromMatrix()` function, applying the following model:

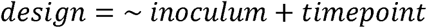

To test genotype effects, we converted the consensus count matrix into the DESeqDataSet while accounting for the genotype effect:

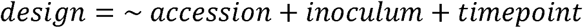

We filtered out low-variance and low-expression genes before performing post hoc tests, which were based on manually defined contrast matrices (e.g., infected vs mock at each time point). We stabilised the variance using the DESeq2-function `lfcShrink(type=”ashr”)` to account for noise and outlier effects. Genes with an absolute log2 fold-change greater than one and an adjusted p-value less than 0.05 were designated as differentially expressed.

### Time course analysis

We performed time-course clustering on focal DEGs for each genotype separately, using a Dirichlet process Gaussian process model (McDowell et al. 2018). This approach addresses three major challenges in RNAseq time-series analysis: selection of the optimal number of clusters, accurate modelling of temporal trajectories and handling uncertainty. Accordingly, a Dirichlet Process is used to dynamically determine the number of clusters based on the input data without *a priori* assumptions. Following this, a Gaussian process is used to model non-random changes in gene expression over time. Smooth expression trajectories are fitted for each cluster, dynamically accounting for intra-cluster variability by including a marginal variance term. Bayesian inference is then used to quantify uncertainty and estimate cluster membership probabilities. We selected clusters using a Maximum a Posteriori (MAP) approach, which accounts for missing or noisy data. This model is implemented in the DP_GP software (https://github.com/PrincetonUniversity/DP_GP_cluster?tab=readme-ov-file, last accessed Feb. 2025).

### Gene Ontology Analysis

To characterise the putative function of gene sets, we extracted gene ontology terms (GO-terms) using the tool PANNZER2 and filtered for a reliability estimate (positive predictive value, PPV) greater 0.4 (Törönen and Holm 2022). Redundant or outdated GO-terms were filtered using REVIGO v1.8.1 implemented in a custom script (Supek et al. 2011). We then performed Gene Set Enrichment Analysis (GSEA) for each genotype at defined timepoints using the clusterprofiler v4.14.4 package with the options `pvalueCutoff=0.05, exponent=0.5, pAdjustMethod=”BH”, by=”fgsea”` (Xu et al. 2024). We conducted the analysis of the REVIGO annotations discretely for molecular functions and biological processes. Putative protein IDs of candidate genes were identified using the blastp function of the UniProt database (The UniProt Consortium 2025).

### Functional Characterization

We performed an OrthoFinder v2.5.5 analysis comparing the *S. pennellii* proteome with *A. thaliana* and *S. lycopersicum* to infer putative gene functions based on sequence homology. Protein names were assigned according to their homology with *A. thaliana* for broadly conserved genes and with *S. lycopersicum* for Solanum-specific genes. Additionally, using a custom pipeline, we previously identified putative receptor-like proteins and receptor-like kinases specifically for *S. pennellii* (see online resources).

### Gene-Regulatory-Network

We used the Plant Transcription Factor Database to predict *S. pennellii* transcription factors based on the proteome sequences (Jin et al. 2017). We then generated a directed gene regulatory network based on variance-stabilised read counts of all inoculated samples using the R-package GENIE3 v1.28.0. For this, we used the `GENIE3()` function. We edge-weight- filtered the network using a ranked inflexion point thresholding approach developed by previous studies (Tominello-Ramirez et al. 2024). We used the same thresholding method to determine hub genes based on eigencentrality (see digital resources).

### Statistical analysis, Visualisation and basic programming

Most analyses were conducted in the R programming environment v4.4.1 within RStudio v2024.12.0. The tidyverse package v2.0.0 was used for data manipulation *dplyr* v1.1.4, visualisation *ggplot2* v3.5.1, and string processing *stringr* v1.5.1. Principal component analysis was performed using the pcatools v2.18.0 package, while ComplexHeatmap v2.22.0 was used for heatmap visualisation. All other plots were generated with *ggplot2*. Vectorised graphics were refined in Inkscape to create multi-panel figures and ensure consistent aesthetics, while GIMP was used for exemplary overlays. For more details about the software versions, please refer to the session info in the online resources.

### Online resources

The scripts and pipelines used in this study are publicly available at github.com/PHYTOPatCAU/Spen_lag_phase_RNAseq. The pipeline for genome annotation improvement is available at github.com/PHYTOPatCAU/SolanumPhylotranscriptomics. The pipeline documenting RLP and RLK prediction is available at github.com/PHYTOPatCAU/RLP_identification

## Declarations

### Ethics approval and consent to participate

Not applicable

### Consent for publication

Not applicable

### Availability of data and materials

All raw files and the final count matrices are publicly available at the NCBI Gene Expression Omnibus (GEO) under the accession GSE292483.

## Funding

This research received partial funding from the DFG (STA1547/6 and STA1547/8) and the ANR (ANR-21-CE20-30) for the ResiDEvo collaborative project.

### Authors’ contributions

SE, LG and TvdL planned and conducted the lab experiments. SE, ATH and NJ developed and integrated the software and analyses used in this study. SE performed data analysis and visualisation. SE and RS conceived the study and wrote the manuscript. All authors reviewed and approved the manuscript.

## Acknowledgements

We would like to acknowledge Susanne Kleingarn for her assistance during the RNA isolation process. Our warmest gratitude also belongs to Hendrik Seide, who supported us with critical discussions.

