## Supplementary Figures for "Temporal Shifts in Gene Expression Drive Quantitative Resistance to a Necrotrophic Fungus in a Tomato Crop Wild Relative"

### Supplementary Material

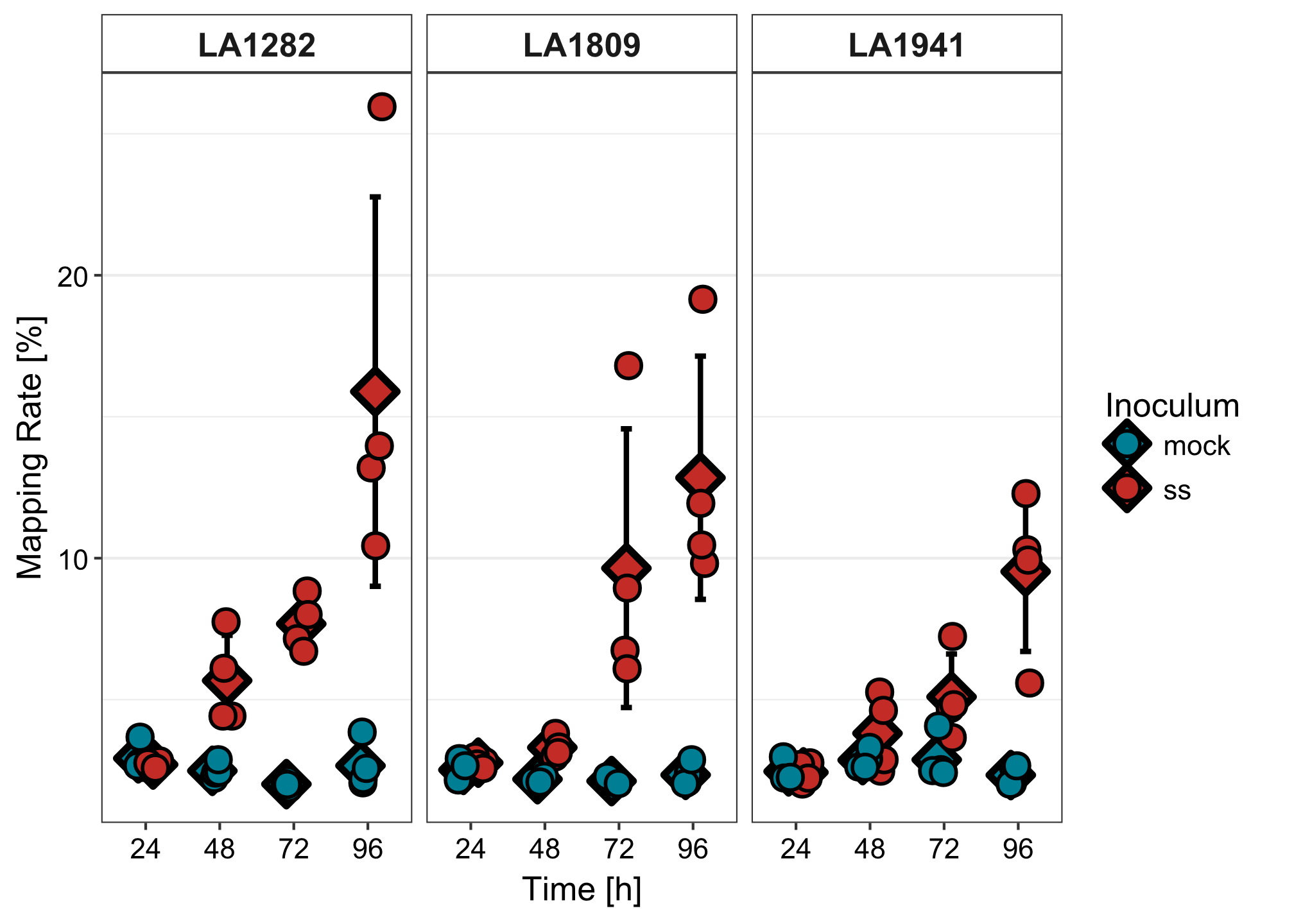

**Suppl. fig. 1: Mapping rate of fungal reads.** Diamonds indicate the mean mapping rate per time point and treatment. Dots represent individual samples, and whiskers depict the standard deviation.

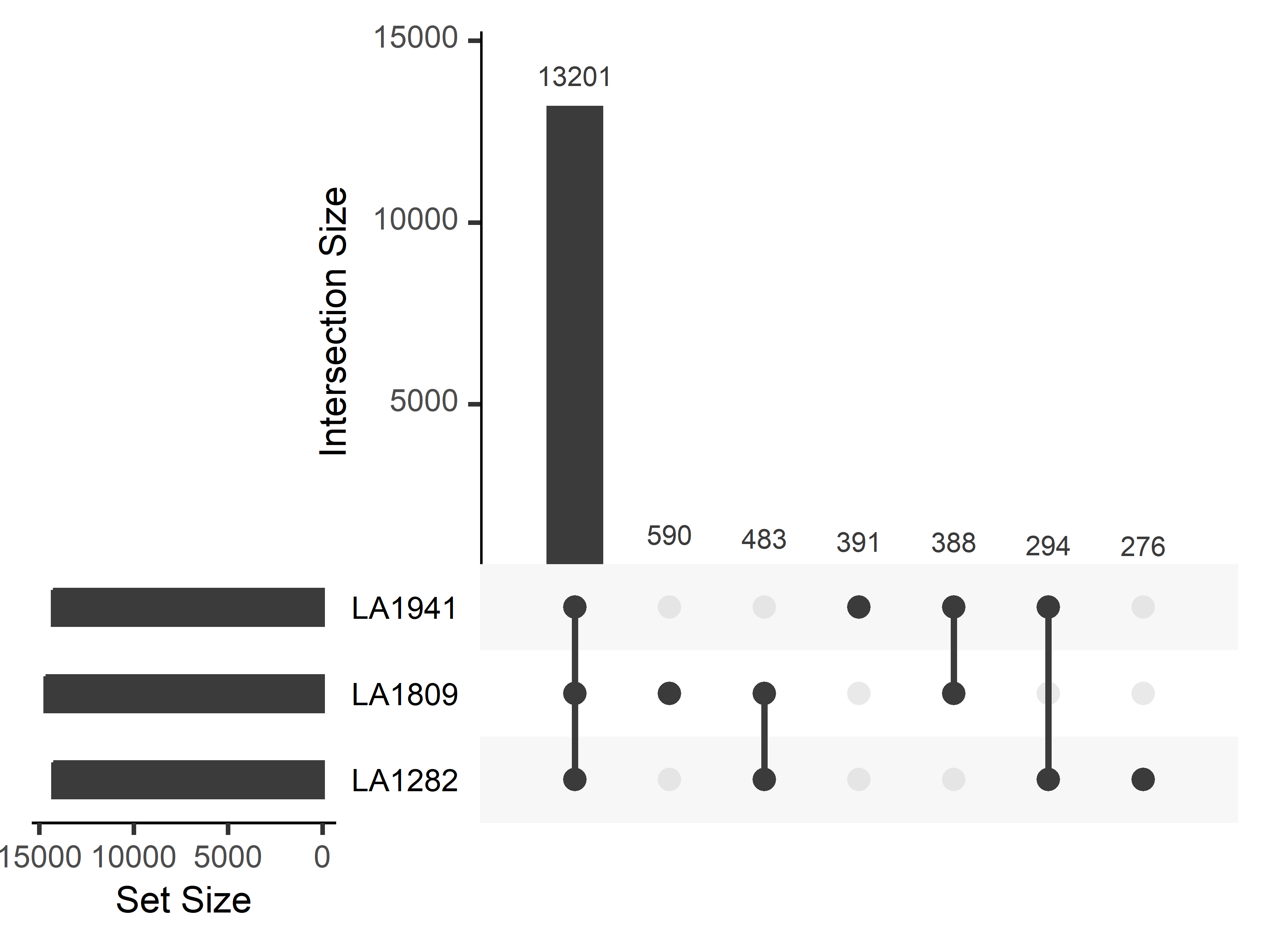

**Suppl. fig. 2: Number of shared and unique expressed genes.**

**Suppl. Table 1 DEGs 48 hpi, LA1282 inf-mock**

| **GeneID** | **L2FC** | **padj** | **Putative Function** |
| --- | --- | --- | --- |
| Sopen08g009740 | 4.296 | 0.0004 | LOX1 |
| Sopen08g008720 | 4.276 | 0.0007 | Sopen08g008720 |
| Sopen08g008720 | 4.276 | 0.0007 | Sopen08g008720 |
| Sopen01g001520 | 4.228 | 0.0039 | NAD(P)-binding Rossmann-fold superfamily protein |
| Sopen02g031830 | 3.959 | 0.0046 | DOX1 |
| Sopen12g023430 | 3.601 | 0.0007 | Sopen12g023430 |
| Sopen10g033140 | 3.365 | 0.0046 | CytochromeP450 (Sopen10g033140) |
| Sopen04g012680 | 3.146 | 0.0123 | Glutamatedecarboxylase |
| Sopen09g028060 | 3.115 | 0.0123 | ERF |
| Sopen04g026840 | 3.044 | 0.0136 | Pathogen-related protein |
| Sopen06g011740 | 3.028 | 0.0080 | Oxalate--CoAligase |
| Sopen01g051340 | 2.927 | 0.0139 | divinylether synthase |
| Sopen09g036010 | 2.868 | 0.0146 | Glycosyltransferase |
| Sopen02g037820 | 2.709 | 0.0122 | CCoAOMT1 (Sopen02g037820) |
| g.plus_peak_18795 | 2.656 | 0.0175 | g.plus_peak_18795 |
| Sopen01g026150 | 2.647 | 0.0080 | beta-1,3-endoglucanase (Sopen01g026150) |
| Sopen10g028890 | 2.614 | 0.0185 | CytochromeP450 (Sopen10g028890) |
| Sopen04g029490 | 2.546 | 0.0099 | DAHPsynthase 2 precursor |
| Sopen03g007540 | 2.509 | 0.0122 | no symbol available (Sopen03g007540) |
| Sopen03g007540 | 2.509 | 0.0122 | no symbol available (Sopen03g007540) |
| Sopen12g023420 | 2.446 | 0.0080 | Sopen12g023420 |
| Sopen02g031030 | 2.432 | 0.0139 | Sopen02g031030 |
| Sopen02g031030 | 2.432 | 0.0139 | Sopen02g031030 |
| Sopen11g027650 | 2.373 | 0.0221 | PUB24 |
| Sopen10g028130 | 2.337 | 0.0122 | Ammoniumtransporter |
| Sopen07g024910 | 2.305 | 0.0038 | EF501822(Sopen07g024910) |
| Sopen01g002940 | 2.292 | 0.0122 | syntaxin-121-like |
| Sopen03g029450 | 2.262 | 0.0185 | Bioticcell death-associated protein |
| Sopen08g023280 | 2.227 | 0.0221 | Polyphenoloxidase |
| Sopen03g041120 | 2.117 | 0.0207 | CytochromeP450 (Sopen03g041120) |
| Sopen09g033940 | 2.040 | 0.0139 | Majorallergen Pru ar.1 (Sopen09g033940) |
| Sopen12g034850 | 2.020 | 0.0267 | Fattyacid desaturase |
| Sopen04g025580 | 2.002 | 0.0080 | Sopen04g025580 |
| Sopen01g026160 | 1.994 | 0.0080 | Sopen01g026160 |
| Sopen12g021540 | 1.909 | 0.0164 | Sopen12g021540 |
| Sopen03g033590 | 1.859 | 0.0221 | ENO1 |
| Sopen12g032770 | 1.858 | 0.0292 | Majorallergen Pru ar.1 (Sopen12g032770) |
| Sopen10g032770 | 1.818 | 0.0330 | AtCWIN2, AtcwINV2, CWINV2 |
| Sopen01g046060 | 1.817 | 0.0221 | Alpha/beta-Hydrolasessuperfamily protein |
| Sopen03g033250 | 1.814 | 0.0122 | Sopen03g033250 |
| Sopen11g009900 | 1.807 | 0.0252 | Peroxidase |
| Sopen07g024490 | 1.800 | 0.0221 | Cytochromeb5.1 |
| Sopen04g028240 | 1.703 | 0.0340 | WRKY51 |
| Sopen10g034420 | 1.686 | 0.0102 | GlutathioneS-transferase |
| Sopen08g028940 | 1.683 | 0.0207 | Serineprotease inhibitor, potato inhibitor I-type family protein |
| Sopen08g022430 | 1.682 | 0.0340 | aromaticamino acid decarboxylase 2 |
| Sopen08g028950 | 1.675 | 0.0090 | OSM34 |
| Sopen08g023580 | 1.631 | 0.0340 | UDP-Glycosyltransferasesuperfamily protein |
| Sopen05g011470 | 1.630 | 0.0292 | WRKY45 |
| Sopen05g004840 | 1.547 | 0.0221 | Glycosyl transferase, family 14 |
| Sopen01g033730 | 1.542 | 0.0357 | Probable magnesium transporter |
| Sopen03g041450 | 1.541 | 0.0355 | RING/U-boxsuperfamily protein |
| Sopen12g034880 | 1.540 | 0.0368 | CER1-L1 |
| Sopen01g048960 | 1.529 | 0.0340 | PR-1 protein 38 (Sopen01g048960) |
| Sopen08g025910 | 1.514 | 0.0207 | Sopen08g025910 |
| Sopen01g040940 | 1.497 | 0.0080 | PR4 (Sopen01g040940) |
| Sopen03g029570 | 1.465 | 0.0389 | Proteinase inhibitor II |
| Sopen05g029280 | 1.457 | 0.0292 | Phosphoglyceratemutase family protein |
| Sopen01g026170 | 1.433 | 0.0102 | beta-1,3-endoglucanase (Sopen01g026170) |
| Sopen09g034580 | 1.415 | 0.0262 | ABCG39 |
| Sopen08g022500 | 1.366 | 0.0221 | Tyraminen-hydroxycinnamoyl transferase |
| Sopen03g028580 | 1.348 | 0.0340 | Phosphatecarrier protein, mitochondrial |
| Sopen02g037830 | 1.331 | 0.0340 | CCoAOMT1 (Sopen02g037830) |
| Sopen10g034470 | 1.310 | 0.0357 | no symbol available (Sopen10g034470) |
| Sopen10g034470 | 1.310 | 0.0357 | no symbol available (Sopen10g034470) |
| Sopen01g044660 | 1.279 | 0.0340 | TPS20 |
| Sopen12g029340 | 1.227 | 0.0500 | Alpha/beta-hydrolasessuperfamily protein |
| Sopen01g048970 | 1.211 | 0.0162 | PR-1 protein 38 (Sopen01g048970) |
| Sopen05g004580 | 1.210 | 0.0207 | Lipidphosphate phosphatase.1 |
| Sopen05g004580 | 1.210 | 0.0207 | Lipidphosphate phosphatase.1 |
| Sopen01g034780 | 1.190 | 0.0364 | Subtilisin-likeprotease |
| Sopen01g040950 | 1.168 | 0.0137 | PR4 (Sopen01g040950) |
| Sopen01g040950 | 1.168 | 0.0137 | PR4 (Sopen01g040950) |
| Sopen07g001220 | 1.123 | 0.0448 | Chitinase |
| Sopen09g006690 | 1.103 | 0.0500 | Sopen09g006690 |
| Sopen07g024880 | 1.031 | 0.0221 | EF501822(Sopen07g024880) |
| Sopen11g001970 | 1.006 | 0.0500 | CMPG1, ATCMPG1 |

**Suppl. Tab. 2: DEGs LA1941, 48 hpi inf vs mock**

| **GeneID** | **L2FC** | **padj** | **funct_unique** |
| --- | --- | --- | --- |
| Sopen03g039350 | 1.43 | 0.003 | Sopen03g039350 |
| Sopen03g040350 | -1.25 | 0.002 | beta-galactosidase3 |
| Sopen04g001120 | 1.81 | 0.000 | MYBD |
| Sopen04g024830 | 1.15 | 0.003 | PLAT3 |
| Sopen04g027010 | 2.02 | 0.001 | EXT3 |
| Sopen04g035160 | 1.81 | 0.001 | Thaumatin-likeprotein.1 |
| Sopen06g024710 | -1.58 | 0.000 | OXS3 |
| Sopen08g022270 | 1.03 | 0.005 | MFP-a |
| Sopen09g004270 | 1.21 | 0.002 | Alpha/beta-Hydrolasessuperfamily protein |
| Sopen09g032860 | 1.05 | 0.003 | ERF1B |
| Sopen10g001910 | 1.22 | 0.001 | FLA2 |

**Suppl. Tab. 3: GSEA LA1809, 48hpi**

| **ID** | **Description** | **NES** | **p.adjust** | **setSize** |
| --- | --- | --- | --- | --- |
| GO:0006412 | translation | 4.050 | 0.0001 | 113 |
| GO:0002181 | cytoplasmic translation | 4.033 | 0.0001 | 24 |
| GO:0002191 | cap-dependent translational initiation | 3.821 | 0.0001 | 25 |
| GO:0000028 | ribosomal small subunit assembly | 3.799 | 0.0001 | 161 |
| GO:0016226 | iron-sulfur cluster assembly | 3.492 | 0.0002 | 22 |
| GO:0006520 | amino acid metabolic process | 3.246 | 0.0010 | 13 |
| GO:0006570 | tyrosine metabolic process | 3.009 | 0.0003 | 52 |
| GO:0006414 | translational elongation | 2.976 | 0.0022 | 32 |
| GO:0006112 | energy reserve metabolic process | 2.888 | 0.0016 | 41 |
| GO:0006734 | NADH metabolic process | 2.842 | 0.0160 | 17 |
| GO:0006457 | protein folding | 2.823 | 0.0001 | 103 |
| GO:0042254 | ribosome biogenesis | 2.809 | 0.0019 | 60 |
| GO:1900864 | mitochondrial RNA modification | 2.792 | 0.0207 | 21 |
| GO:0006760 | folic acid-containing compound metabolic process | 2.676 | 0.0083 | 45 |
| GO:0032988 | protein-RNA complex disassembly | 2.596 | 0.0307 | 29 |
| GO:0006913 | nucleocytoplasmic transport | 2.438 | 0.0092 | 74 |
| GO:0006227 | dUDP biosynthetic process | 2.324 | 0.0470 | 60 |

**Suppl. Tab. 4: GSEA LA1941, 48hpi**

| **ID** | **Description** | **NES** | **p.adjust** | **setSize** |
| --- | --- | --- | --- | --- |
| GO:0002181 | cytoplasmic translation | 3.238 | 0.026 | 21 |
| GO:0007018 | microtubule-based movement | 3.104 | 0.026 | 16 |
| GO:0006952 | defense response | 2.928 | 0.008 | 83 |

**Suppl. Tab. 5: cRLP-Genes with differential expression between LA1282 and LA1809 in basal conditions (24hpi mock).**

| **GeneID** | **log2FoldChange** | **padj** |
| --- | --- | --- |
| Sopen01g002540 | 2.425 | 0.0110 |
| Sopen06g003150 | 2.095 | 0.0002 |
| Sopen01g001720 | 1.466 | 0.0180 |
| Sopen05g005800 | -1.113 | 0.0135 |
| Sopen01g001870 | -1.266 | 0.0247 |
| Sopen01g042220 | -2.552 | 0.0103 |
| Sopen06g014090 | -4.597 | 0.0000 |

**Suppl. Tab. 6: RLK-Genes with differential expression between LA1282 and LA1809 in basal conditions (24hpi mock).**

| **GeneID** | **baseMean** | **log2FoldChange** | **padj** |
| --- | --- | --- | --- |
| Sopen02g020810 | 17.1486 | 3.8109 | 0.0000 |
| Sopen06g002020 | 9.8637 | 2.4009 | 0.0000 |
| Sopen01g048860 | 19.3616 | 2.2068 | 0.0010 |
| Sopen12g032540 | 9.8612 | 1.8516 | 0.0017 |
| Sopen01g028710 | 9.1444 | 1.7339 | 0.0076 |
| Sopen08g019890 | 69.3694 | 1.5467 | 0.0002 |
| Sopen01g047490 | 94.4411 | 1.1532 | 0.0002 |
| Sopen03g030900 | 32.0974 | 1.0801 | 0.0128 |
| Sopen05g009260 | 10.0071 | -1.0246 | 0.0073 |
| Sopen05g030910 | 9.1162 | -1.1905 | 0.0042 |
| Sopen02g036420 | 67.9849 | -1.2143 | 0.0000 |
| Sopen06g030490 | 4.3565 | -1.3392 | 0.0249 |
| Sopen08g020310 | 27.3608 | -1.3694 | 0.0003 |
| Sopen10g029820 | 3.9187 | -1.5135 | 0.0135 |
| Sopen02g037670 | 1.9612 | -1.5981 | 0.0368 |
| Sopen04g003810 | 3.5426 | -1.6662 | 0.0184 |
| Sopen03g001960 | 5.2876 | -1.9929 | 0.0078 |
| Sopen08g017000 | 10.9703 | -2.3844 | 0.0002 |
| Sopen11g001980 | 13.4983 | -2.5488 | 0.0001 |
| Sopen08g020090 | 3.0906 | -2.6146 | 0.0079 |

**Suppl. Tab. 7: PTI-Genes with differential expression between LA1282 and LA1809 in basal conditions (24hpi mock).**

| **gene** | **geneID** | **baseMean** | **L2FC** | **padj** |
| --- | --- | --- | --- | --- |
| cerk1.3 | Sopen07g025160 | 62.3909767014575 | 1.091 | 0.000 |
| bak.1 | Sopen01g047490 | 94.4411297319458 | 1.153 | 0.000 |

**Suppl. Tab. 8: Edgeweight between all focal DEGs and the GRN hub WRKY6**

| **regulatoryGene** | **targetGene** | **weight** |
| --- | --- | --- |
| Sopen02g011350 | Sopen08g009740 | 0.028360696 |
| Sopen02g011350 | Sopen01g026150 | 0.028051527 |
| Sopen02g011350 | Sopen08g023280 | 0.026719102 |
| Sopen02g011350 | Sopen02g031830 | 0.02569113 |
| Sopen02g011350 | Sopen12g021540 | 0.025036 |
| Sopen02g011350 | Sopen01g044660 | 0.023434218 |
| Sopen02g011350 | Sopen08g023580 | 0.023399628 |
| Sopen02g011350 | Sopen01g046060 | 0.022046422 |
| Sopen02g011350 | Sopen07g024910 | 0.021682299 |
| Sopen02g011350 | Sopen04g025580 | 0.021600303 |
| Sopen02g011350 | Sopen05g029280 | 0.021274326 |
| Sopen02g011350 | Sopen03g033250 | 0.020913354 |
| Sopen02g011350 | Sopen03g041450 | 0.020645404 |
| Sopen02g011350 | Sopen10g034420 | 0.019906255 |
| Sopen02g011350 | Sopen04g029490 | 0.019795622 |
| Sopen02g011350 | Sopen12g034850 | 0.019026858 |
| Sopen02g011350 | Sopen07g024880 | 0.018282362 |
| Sopen02g011350 | Sopen03g028580 | 0.01742883 |
| Sopen02g011350 | Sopen09g036010 | 0.017111209 |
| Sopen02g011350 | Sopen12g032770 | 0.016996095 |
| Sopen02g011350 | Sopen03g029570 | 0.016597635 |
| Sopen02g011350 | Sopen01g040950 | 0.016414153 |
| Sopen02g011350 | Sopen01g001520 | 0.016053445 |
| Sopen02g011350 | Sopen08g008720 | 0.015724586 |
| Sopen02g011350 | Sopen01g048960 | 0.015687376 |
| Sopen02g011350 | Sopen01g040940 | 0.015633476 |
| Sopen02g011350 | Sopen05g004840 | 0.015351446 |
| Sopen02g011350 | Sopen01g034780 | 0.014946166 |
| Sopen02g011350 | Sopen08g022500 | 0.014828637 |
| Sopen02g011350 | Sopen01g033730 | 0.014503793 |
| Sopen02g011350 | Sopen04g012680 | 0.013928743 |
| Sopen02g011350 | Sopen03g007540 | 0.013856238 |
| Sopen02g011350 | Sopen09g028060 | 0.013665484 |
| Sopen02g011350 | Sopen01g026170 | 0.0136052 |
| Sopen02g011350 | Sopen01g026160 | 0.01340945 |
| Sopen02g011350 | Sopen08g028950 | 0.013371735 |
| Sopen02g011350 | Sopen07g024490 | 0.012750647 |
| Sopen02g011350 | Sopen04g026840 | 0.012431308 |
| Sopen02g011350 | Sopen02g031030 | 0.012424489 |
| Sopen02g011350 | Sopen04g028240 | 0.012364526 |
| Sopen02g011350 | Sopen01g048970 | 0.012256411 |
| Sopen02g011350 | Sopen07g001220 | 0.012173157 |
| Sopen02g011350 | Sopen01g002940 | 0.012161811 |
| Sopen02g011350 | Sopen08g025910 | 0.012129958 |
| Sopen02g011350 | Sopen11g027650 | 0.012106668 |
| Sopen02g011350 | Sopen05g011470 | 0.011980095 |
| Sopen02g011350 | Sopen03g033590 | 0.011918254 |
| Sopen02g011350 | Sopen12g034880 | 0.011818788 |
| Sopen02g011350 | Sopen02g037830 | 0.011766374 |
| Sopen02g011350 | Sopen10g028130 | 0.011642719 |
| Sopen02g011350 | Sopen02g037820 | 0.011336435 |
| Sopen02g011350 | Sopen01g051340 | 0.0109734 |
| Sopen02g011350 | Sopen10g033140 | 0.010961717 |
| Sopen02g011350 | Sopen10g032770 | 0.010744659 |
| Sopen02g011350 | Sopen09g034580 | 0.010057291 |
| Sopen02g011350 | Sopen06g011740 | 0.010051028 |
| Sopen02g011350 | g.plus_peak_18795 | 0.009756631 |
| Sopen02g011350 | Sopen11g009900 | 0.009656683 |
| Sopen02g011350 | Sopen09g033940 | 0.008887826 |
| Sopen02g011350 | Sopen08g022430 | 0.008757446 |
| Sopen02g011350 | Sopen03g029450 | 0.008587933 |
| Sopen02g011350 | Sopen12g029340 | 0.008311617 |
| Sopen02g011350 | Sopen12g023430 | 0.008074835 |
| Sopen02g011350 | Sopen10g028890 | 0.007815482 |
| Sopen02g011350 | Sopen12g023420 | 0.007661805 |
| Sopen02g011350 | Sopen05g004580 | 0.007359228 |
| Sopen02g011350 | Sopen11g001970 | 0.006641716 |
| Sopen02g011350 | Sopen03g041120 | 0.006346855 |
| Sopen02g011350 | Sopen08g028940 | 0.005798289 |
| Sopen02g011350 | Sopen10g034470 | 0.005191843 |

**Suppl. Tab. 9: Focal Degs basal Expression 24 hpi mock, LA1282 vs. LA1809**

| **GeneID** | **L2FC** | **padj** | **DEG** | **funct_unique** |
| --- | --- | --- | --- | --- |
| Sopen01g048970 | 7.285 | 2.848E-31 | DEG | PR-1 protein 38 (Sopen01g048970) |
| Sopen03g029570 | 7.154 | 1.463E-11 | DEG | Proteinase inhibitor II |
| Sopen08g028950 | 6.800 | 1.443E-19 | DEG | OSM34 |
| Sopen01g026150 | 5.878 | 3.160E-07 | DEG | beta-1,3-endoglucanase (Sopen01g026150) |
| Sopen01g026170 | 5.835 | 1.093E-20 | DEG | beta-1,3-endoglucanase (Sopen01g026170) |
| Sopen01g048960 | 5.388 | 1.213E-05 | DEG | PR-1 protein 38 (Sopen01g048960) |
| Sopen08g009740 | 5.324 | 4.281E-06 | DEG | LOX1 |
| Sopen01g026160 | 5.253 | 8.686E-12 | DEG | Sopen01g026160 |
| Sopen12g034850 | 5.250 | 3.094E-05 | DEG | Fattyacid desaturase |
| Sopen08g023280 | 5.203 | 1.094E-05 | DEG | Polyphenoloxidase |
| Sopen03g033250 | 5.034 | 2.125E-11 | DEG | Sopen03g033250 |
| Sopen02g031830 | 4.470 | 9.971E-05 | DEG | DOX1 |
| Sopen04g025580 | 4.386 | 4.013E-08 | DEG | Sopen04g025580 |
| Sopen10g028890 | 4.361 | 1.105E-04 | DEG | CytochromeP450 (Sopen10g028890) |
| Sopen08g028940 | 4.248 | 2.884E-06 | DEG | Serineprotease inhibitor, potato inhibitor I-type family protein |
| Sopen01g040940 | 4.049 | 6.416E-13 | DEG | PR4 (Sopen01g040940) |
| Sopen07g024490 | 3.691 | 7.150E-05 | DEG | Cytochromeb5.1 |
| Sopen09g034580 | 3.608 | 1.487E-06 | DEG | ABCG39 |
| Sopen03g028580 | 3.591 | 1.924E-05 | DEG | Phosphatecarrier protein, mitochondrial |
| Sopen02g031030 | 3.439 | 8.870E-04 | DEG | Sopen02g031030 |
| Sopen10g034420 | 3.423 | 6.974E-09 | DEG | GlutathioneS-transferase |
| Sopen01g034780 | 3.376 | 5.176E-07 | DEG | Subtilisin-likeprotease |
| Sopen01g040950 | 3.276 | 8.685E-13 | DEG | PR4 (Sopen01g040950) |
| Sopen01g051340 | 3.123 | 5.724E-03 | DEG | divinylether synthase |
| Sopen09g028060 | 3.104 | 4.198E-03 | DEG | ERF |
| Sopen12g029340 | 3.090 | 9.639E-04 | DEG | Alpha/beta-hydrolasessuperfamily protein |
| Sopen03g029450 | 3.030 | 1.547E-03 | DEG | Bioticcell death-associated protein |
| Sopen05g011470 | 3.013 | 2.269E-04 | DEG | WRKY45 |
| Sopen12g034880 | 2.956 | 2.449E-03 | DEG | CER1-L1 |
| Sopen10g032770 | 2.791 | 2.245E-03 | DEG | AtCWIN2, AtcwINV2, CWINV2 |
| Sopen08g008720 | 2.696 | 1.478E-03 | DEG | Sopen08g008720 |
| Sopen01g046060 | 2.692 | 1.659E-04 | DEG | Alpha/beta-Hydrolasessuperfamily protein |
| Sopen04g026840 | 2.656 | 5.135E-03 | DEG | Pathogen-relatedprotein |
| Sopen08g022430 | 2.564 | 6.495E-03 | DEG | aromaticamino acid decarboxylase 2 |
| Sopen11g009900 | 2.518 | 1.355E-03 | DEG | Peroxidase |
| Sopen02g037830 | 2.410 | 7.291E-04 | DEG | CCoAOMT1 (Sopen02g037830) |
| Sopen01g002940 | 2.301 | 2.944E-03 | DEG | syntaxin-121-like |
| Sopen09g036010 | 2.281 | 1.074E-02 | DEG | Glycosyltransferase |
| Sopen04g012680 | 2.205 | 7.548E-03 | DEG | Glutamatedecarboxylase |
| Sopen10g033140 | 2.193 | 7.211E-03 | DEG | CytochromeP450 (Sopen10g033140) |
| Sopen06g011740 | 2.179 | 4.220E-03 | DEG | Oxalate--CoAligase |
| Sopen10g028130 | 2.160 | 2.367E-03 | DEG | Ammoniumtransporter |
| Sopen05g004580 | 2.160 | 7.142E-07 | DEG | Lipidphosphate phosphatase.1 |
| Sopen03g041450 | 2.101 | 1.167E-02 | DEG | RING/U-boxsuperfamily protein |
| Sopen03g007540 | 1.859 | 1.741E-02 | DEG | no symbol available (Sopen03g007540) |
| Sopen12g032770 | 1.799 | 1.221E-02 | DEG | Majorallergen Pru ar.1 (Sopen12g032770) |
| Sopen08g022500 | 1.782 | 6.362E-03 | DEG | Tyraminen-hydroxycinnamoyl transferase |
| Sopen02g037820 | 1.769 | 1.656E-02 | DEG | CCoAOMT1 (Sopen02g037820) |
| Sopen11g001970 | 1.723 | 1.792E-02 | DEG | CMPG1, ATCMPG1 |
| Sopen12g021540 | 1.604 | 7.014E-03 | DEG | Sopen12g021540 |
| Sopen01g044660 | 1.589 | 4.343E-02 | DEG | TPS20 |
| Sopen07g001220 | 1.513 | 2.222E-02 | DEG | Chitinase |
| Sopen12g023420 | 1.437 | 1.756E-02 | DEG | NA |
| Sopen09g006690 | 1.420 | 2.016E-02 | DEG | Sopen09g006690 |
| Sopen04g029490 | 1.343 | 1.350E-02 | DEG | DAHPsynthase 2 precursor |
| Sopen07g024880 | 1.266 | 1.743E-03 | DEG | EF501822(Sopen07g024880) |
| Sopen11g027650 | 1.185 | 5.776E-02 | Not DEG | PUB24 |
| Sopen04g028240 | 1.166 | 6.052E-02 | Not DEG | WRKY51 |
| Sopen03g041120 | 1.001 | 5.923E-02 | Not DEG | CytochromeP450 (Sopen03g041120) |
| Sopen10g034470 | 0.998 | 4.959E-02 | Not DEG | no symbol available (Sopen10g034470) |
| Sopen05g004840 | 0.912 | 8.509E-02 | Not DEG | Glycosyl transferase, family 14 |
| Sopen01g033730 | 0.854 | 1.083E-01 | Not DEG | Probable magnesium transporter |
| g.plus_peak_18795 | 0.845 | 1.553E-01 | Not DEG | g.plus_peak_18795 |
| Sopen09g033940 | 0.825 | 9.174E-02 | Not DEG | Majorallergen Pru ar.1 (Sopen09g033940) |
| Sopen08g025910 | 0.745 | 8.043E-02 | Not DEG | Sopen08g025910 |
| Sopen12g023430 | 0.667 | 1.580E-01 | Not DEG | NA |
| Sopen01g001520 | 0.660 | 2.302E-01 | Not DEG | NAD(P)-bindingRossmann-fold superfamily protein |
| Sopen07g024910 | 0.570 | 2.105E-01 | Not DEG | EF501822(Sopen07g024910) |
| Sopen08g023580 | 0.569 | 2.232E-01 | Not DEG | UDP-Glycosyltransferasesuperfamily protein |
| Sopen03g033590 | 0.421 | 3.939E-01 | Not DEG | ENO1 |
| Sopen05g029280 | 0.025 | 1.000E+00 | Not DEG | Phosphoglyceratemutase family protein |
